# Lipid Droplet Proteome Reveals that Associated ATGL and Anchored Mitochondria Lead to Higher Skeletal Muscle Insulin Sensitivity in Endurance Athletes than Type 2 Diabetes Mellitus

**DOI:** 10.1101/2025.07.15.665019

**Authors:** Xiaochuan Fu, Zhihua Zhang, Lijun Jiang, Qiumin Liao, Yilu Xu, Yang Wang, Zhen Cao, Jifeng Wang, Xiaojuan Wei, Yan Gao, Yue Li, Haijun Wang, Yin Pei, Fan Hu, Shuyan Zhang, Xin Zhang, Hongchao Zhang, Pingsheng Liu

## Abstract

As the key organelle governed lipid homeostasis, lipid droplets (LDs) in skeletal muscle plays a critical role in regulating systemic insulin sensitivity. In type 2 diabetes mellitus (T2DM) patients intramyocellular lipid (IMCL) content negatively associates with insulin sensitivity, while endurance athletes exhibit IMCL levels were positively correlated to insulin sensitivity, which is known as the athlete’s paradox. To solve this paradox, LDs were isolated from skeletal muscle samples of seven athletes and nine T2DM patients, and the first proteome dataset of human skeletal muscle LD was established. Comparative proteomic analysis revealed 733 upregulated proteins in athlete LDs, primarily enriched in mitochondria, and 755 downregulated proteins, mainly associated with cytoskeletal components, relative to T2DM patients. Integrated analysis with tissue proteomics further indicated enhanced energy metabolism activity of LD-anchored mitochondria (LDAM) in athletes. Notably, adipose triglyceride lipase (ATGL) was specifically upregulated by 2.97-fold on LDs in athletes, while its overall tissue expression remained unchanged. Moreover, deletion of ATGL in myoblasts significantly reduced insulin-stimulated AKT phosphorylation. Additionally, skeletal muscle LDs from db/db and ob/ob mice exhibited reduced levels of mitochondrial proteins and ATGL. Together, our findings reveal a LD-based solution for the athlete’s paradox, and identify ATGL as a key regulator of skeletal muscle insulin sensitivity. The results also suggest that the “bad LDs” in the skeletal muscle of T2DM patients are characterized by higher levels of cytoskeletal proteins, reduced LDAM, and lower ATGL compared to the “good LDs” observed in athletes.

## Introduction

Diabetes mellitus (DM) has emerged as one of the fastest-growing global health emergencies of the 21^st^ century, posing significant threats to global health and substantially compromising quality of life^1,2^. The IDF Diabetes Atlas reports that in 2024, approximately 589 million people have diabetes, and this number is projected to reach 853 million by 2050. DM is primarily classified into three types: type 1 diabetes mellitus (T1DM), type 2 diabetes mellitus (T2DM), and gestational diabetes mellitus (GDM)^3^. Among these, T2DM is by far the most prevalent form of the disease, comprising more than 90% of all diabetes cases globally^4^. T2DM is fundamentally characterized by impaired insulin sensitivity in target tissues, including the liver, skeletal muscle, and adipose tissue^5–7^. Skeletal muscle accounts for over 70% of postprandial glucose disposal, and its insulin resistance is considered a pivotal etiological factor in the development of T2DM^5,8–11^.

It is well established that lipid metabolism in skeletal muscle is closely associated with insulin sensitivity. Lipid droplets (LDs), the primary sites for cellular neutral lipid storage, are now recognized as dynamic organelles that play central roles in cellular lipid storage and metabolic regulation^12–17^. Previous studies have shown that LDs exist within skeletal muscle fibers and that the intramyocellular lipid (IMCL) storage negatively associates with insulin sensitivity^18–25^. Paradoxically, endurance-trained athletes exhibit elevated IMCL levels while maintaining higher insulin sensitivity compared to sedentary individuals. This phenomenon is known as the athlete’s paradox^26–31^. This paradox can be partially explained by the increased proportion of type I muscle fibers in endurance athletes, which contain approximately 2.8-fold higher lipid content than type II fibers while exhibiting greater insulin sensitivity^29^. In contrast, individuals with T2DM tend to have a reduced proportion of type I fibers. LDs in skeletal muscle exist in two primary regions: subsarcolemmal (SS) LDs and intermyofibrillar (IMF) LDs^32,33^. T2DM patients predominantly accumulate larger LDs in the SS region of type II fibers, whereas endurance-trained individuals store lipids in a higher number of LDs within the IMF region of type I fibers^34^. During exhaustive exercise, IMF LDs, but not those SS LDs, are utilized for energy supply^35^. Additionally, in the trained state, IMCL lipolysis is enhanced, leading to greater free fatty acid utilization^36,37^. Therefore, skeletal muscle insulin sensitivity is influenced not solely by the total IMCL content but also by the physiological state of LDs.

Accumulated studies have revealed that the physical contact between LDs and mitochondria exists in multiple tissues and plays an essential role in cellular lipid metabolism and energy homeostasis^38–41^. Two types of LD-mitochondria interactions have been described: “kiss-and-run” contacts and anchoring of LDs and mitochondria^42^. LD-anchored mitochondria (LDAM) are identified in oxidative tissues including skeletal muscle, cardiac muscle, and brown adipose tissue (BAT)^38–44^. Recent studies have shown increased LD-mitochondria contact in the skeletal muscle of elite male endurance athletes^45^. Such contacts facilitate direct fatty acid transfer from LDs to mitochondria, enhancing the efficiency of mitochondrial β-oxidation^46–50^. However, morphological methods alone cannot distinguish between dynamic and anchored contacts, highlighting the need for studies investigating LDAM in skeletal muscles with different insulin sensitivities to better understand the relationship between lipid metabolism and insulin signaling.

While mass spectrometry-based proteomics has characterized the skeletal muscle proteome in T2DM patients^51,52^, the absence of LD proteomic data from human skeletal muscle has hindered understanding of their role in insulin resistance and delayed therapeutic development. Despite extensive morphological studies linking LD characteristics to insulin sensitivity^32,34,35^, the functional role of LD-associated proteins in regulating insulin sensitivity remains unclear. As central hubs of neutral lipid metabolism, LD-associated proteins, including triglyceride lipases such as adipose triglyceride lipase (ATGL) and hormone-sensitive lipase (HSL), are critical for maintaining lipid homeostasis and have been found associated with systemic insulin sensitivity^46,47,53–58^. However, no studies have directly examined how the proteomic composition of LDs in human skeletal muscle contributes to insulin sensitivity, leaving a critical gap in our understanding.

To address these gaps, we profiled the proteome of LDs isolated from skeletal muscle of seven athletes and nine T2DM patients, establishing the first human skeletal muscle LD proteome. Athlete LDs showed enrichment of mitochondrial proteins and enhanced LDAM energy metabolism, while ATGL was specifically upregulated on LDs without changes at the tissue level. Functionally, ATGL deletion reduced insulin-stimulated AKT phosphorylation in myoblasts. Together, these findings reveal distinct LD features linked to insulin sensitivity, offer a LD-based perspective on the athlete’s paradox, and identify ATGL as a key regulator of skeletal muscle insulin signaling. Moreover, these findings reveal distinct characteristics of “bad LDs” in T2DM patients and “good LDs” in athletes. These findings advance our understanding of insulin signaling and potential therapeutic strategies in metabolic diseases.

## Results

### Isolation and characterization of LDs from human skeletal muscle

To investigate the characteristic of LDs from human skeletal muscle, we collected fresh skeletal muscle samples from endurance-trained athletes and T2DM patients, with detailed donor information provided in Table 1. Histological and morphological analyses confirmed the typical structural features of skeletal muscle (Fig. S1A-B). Neutral lipids within skeletal muscle cells were visualized using Oil Red O staining (Fig. S1A-B), while transmission electron microscopy (TEM) confirmed the presence of LDs in the tissue and revealed their close spatial proximity to mitochondria, suggesting potential functional interactions (Fig. S1C).

**Table 1.** Clinical characteristics of human participants.

| Characteristic | Athlete | T2DM |
| --- | --- | --- |
|  | (n=7) | (n=9) |
| Age (years) | 20±4 | 60±10 |
| Height (cm) | 175±8 | 170±5 |
| Weight (kg) | 66±12 | 80±12 |
| BMI (kg/m<sup>2</sup>) | 21.37±2.56 | 27.52±3.93 |
| Fasting blood parameters |  |  |
| Glucose (mM) | 4.6±0.9 | 7.9±2.4* |
| TAG (mM) | 0.65±0.26 | 1.92±0.83 |
| Cholesterol (mM) | 3.89±1.05 | 3.56±0.65 |
| HDL (mM) | 1.26±0.24 | 1.01±0.28 |
| LDL (mM) | 2.27±0.79 | 1.99±0.71 |

Values expressed are mean ±SD. BMI, body mass index; TAG, Triacylglycerol; HDL, high-density lipoprotein; LDL, low-density lipoprotein; \*, The glucose of T2DM patients are controlled by medication.

Obtaining high-quality LD fractions is essential for studying LD composition. However, human skeletal muscle, which is rich in connective tissue and intermuscular fat, often yields low LD recovery and is susceptible to contamination from other cellular components. Through the systematic optimization of the previous method^59^, we developed a robust method to isolate and purify high-quality LDs from human skeletal muscle (Fig. 1A). Briefly, connective tissue was carefully removed, and the muscle was minced and homogenized using a frosted Dounce homogenizer, followed by ultracentrifugation to collect the LD fraction (Fig. 1A). The quality of the LDs was first examined by fluorescence microscopy using LipidTOX Red staining (Fig. 1Ba). Isolated LDs exhibited spherical structures and could be stained by the neutral lipid dye LipidTOX Red, with fluorescent signals merging well with brightfield images. Electron microscopic observation revealed partial membrane structures surrounding LDs, suggesting interactions between LDs and other cellular membranes, indicative of higher metabolic activity in skeletal muscle (Fig. 1Ba). Particle size analysis revealed that purified LDs exhibited a normal size distribution, while thin-layer chromatography (TLC) confirmed that purified LDs contained predominantly neutral lipids triacylglycerol (TAG) with minimal phospholipid contamination (Fig. 1Bb-d).

**Figure 1.**
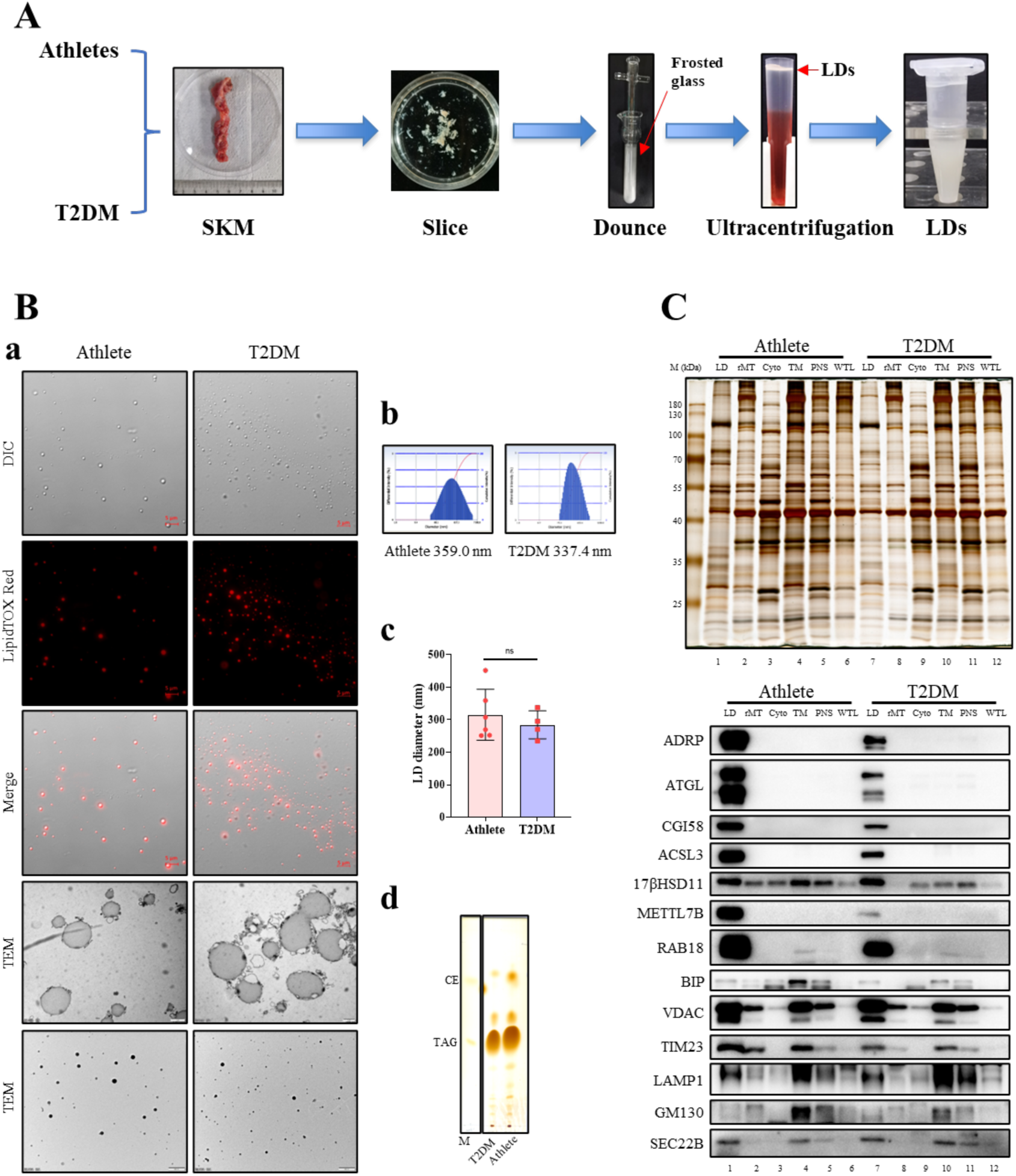
Isolation and characterization of LDs from skeletal muscle of athletes and T2DM patients. (A) The flow chart of LDs isolation from human skeletal muscle. The experimental workflow, as illustrated in the flowchart, consists of five sequential steps: (1) SKM collection, (2) SKM fragmentation, (3) tissue homogenization, (4) homogenate ultracentrifugation, and (5) LDs collection and subsequent analysis. (B) LDs isolated from skeletal muscle of athletes and T2DM patients were stained with LipidTOX Red for 20 minutes and observed by fluorescence microscope (a). Scale bar = 5 µm. The purified LDs were prepared for electron microscopy using two distinct approaches (a). One aliquot was processed through fixation, dehydration, embedding, thin-sectioning, and staining for TEM observation (Scale bar = 1 µm). Another aliquot was directly placed on a Formvar-carbon coated copper grid and observed after positive staining (Scale bar = 500 nm). LDs isolated from skeletal muscle of athletes and T2DM patients were analyzed by a Delsa Nano C particle analyzer (b). The LD diameter statistical results are shown in c. Lipids from LDs were analyzed by thin-layer chromatography (TLC), using cholesteryl ester (CE) and triacylglycerol (TAG) as standards (d). (C) Protein fractions isolated from human skeletal muscle, including lipid droplets (LDs), rough mitochondria (rMT), cytosol (Cyto), total membrane (TM), postnuclear supernatant (PNS), and whole tissue lysate (WTL), were subjected to silver staining and analyzed by Western blot.

To further assess the quality of the isolated LDs, equal amounts of proteins from LD, rough mitochondria (rMT), cytosol (Cyto), total membrane (TM), postnuclear supernatant (PNS), and whole tissue lysate (WTL) fractions were separated by SDS/PAGE. Silver staining showed that LDs displayed a distinct protein profile compared to other fractions (Fig. 1C), indicating the specificity of LD proteins. Western blotting analysis detected seven LD marker proteins (ADRP/PLIN2, ATGL, CGI58, ACSL3, 17βHSD11, METTL7B, and RAB18), an ER marker (BIP), two mitochondrial markers (VDAC and TIM23), a lysosomal marker (LAMP1), a Golgi marker (GM130), and a vesicle marker (SEC22B) (Fig. 1C). All LD-specific marker proteins were highly enriched in the LD fraction. The LD fraction was also enriched in the mitochondrial proteins VDAC and TIM23, while containing lower levels of Golgi and lysosomal proteins, GM130 and LAMP1, respectively.

These findings confirm that the isolated LD preparations are of high purity and suggest that LDs in skeletal muscle exhibit strong interactions with other organelles, particularly mitochondria. The presence of mitochondrial proteins in the LD fraction is consistent with previous reports that LDs functionally interact with mitochondria, particularly in oxidative tissues^38^.

### Proteomic analysis of human skeletal muscle LDs

Using the established purification method, we isolated LDs from skeletal muscle samples of seven athletes and nine T2DM patients and performed proteomic analysis. We also compared the human skeletal muscle LD proteome with previously reported LD proteomes from human liver, human adrenal gland, mouse testis, mouse BAT, and rat heart (Table 2). Among the major LD-associated proteins, all PLIN family members were identified in human skeletal muscle LDs, with PLIN5 detected exclusively in skeletal muscle LDs and absent in human liver and adrenal gland LDs. METTL7B is exclusively found in human skeletal muscle and adrenal gland LDs, with no presence in LDs from other tissues. Within the HSD family proteins, HSD17B13 is uniquely present in human liver LDs, while HSDL1 is only identified in human and mouse skeletal muscle LDs. Regarding the RAB family proteins, 25 RABs were identified in human skeletal muscle LDs, which is consistent with 26 RABs found previously in mouse skeletal muscle LDs^43^. But only RAB6C, RAB9B, and RAB33B were specifically detected in human skeletal muscle LDs, whereas RAB4B is solely identified in skeletal muscle.

**Table 2.**
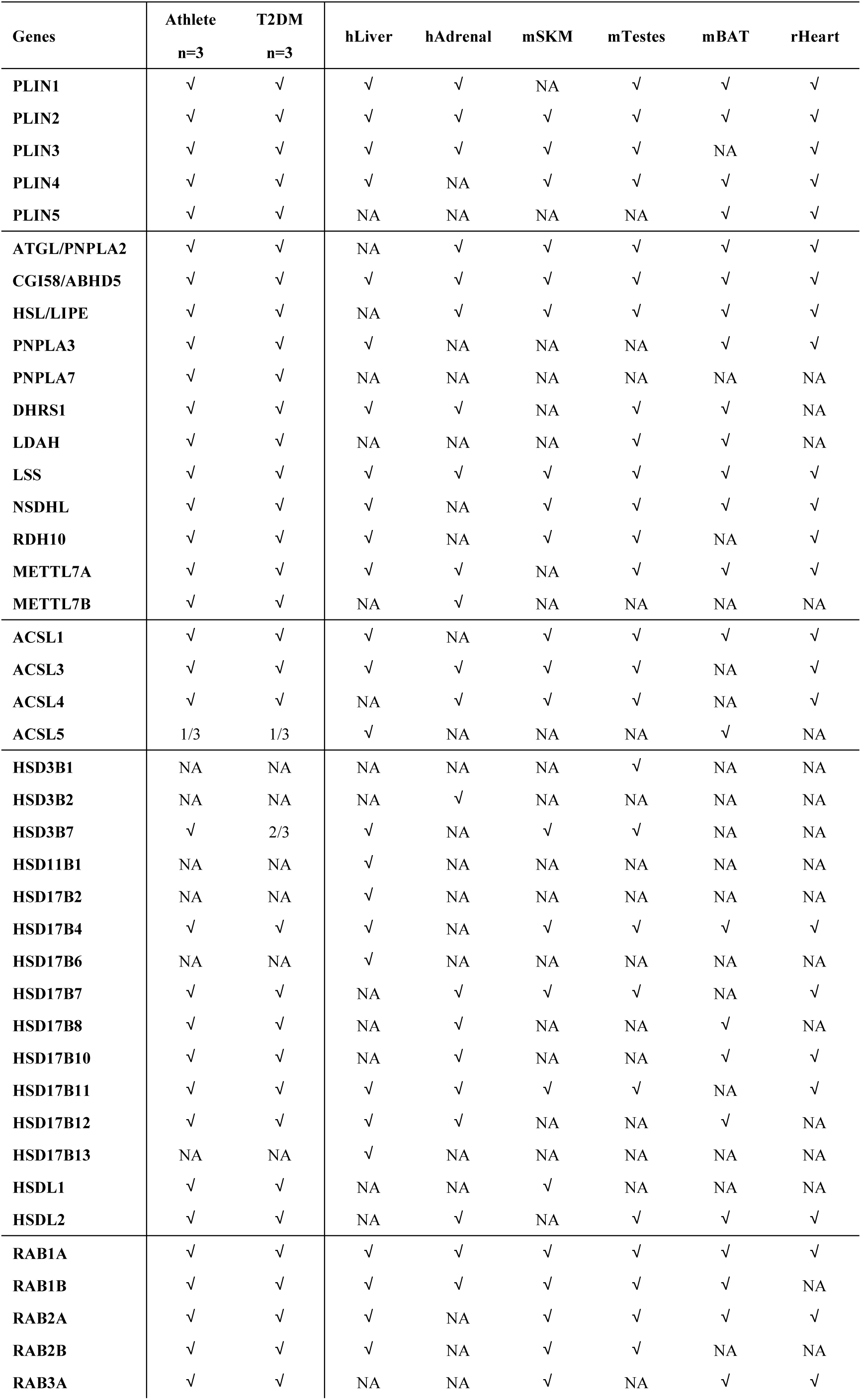
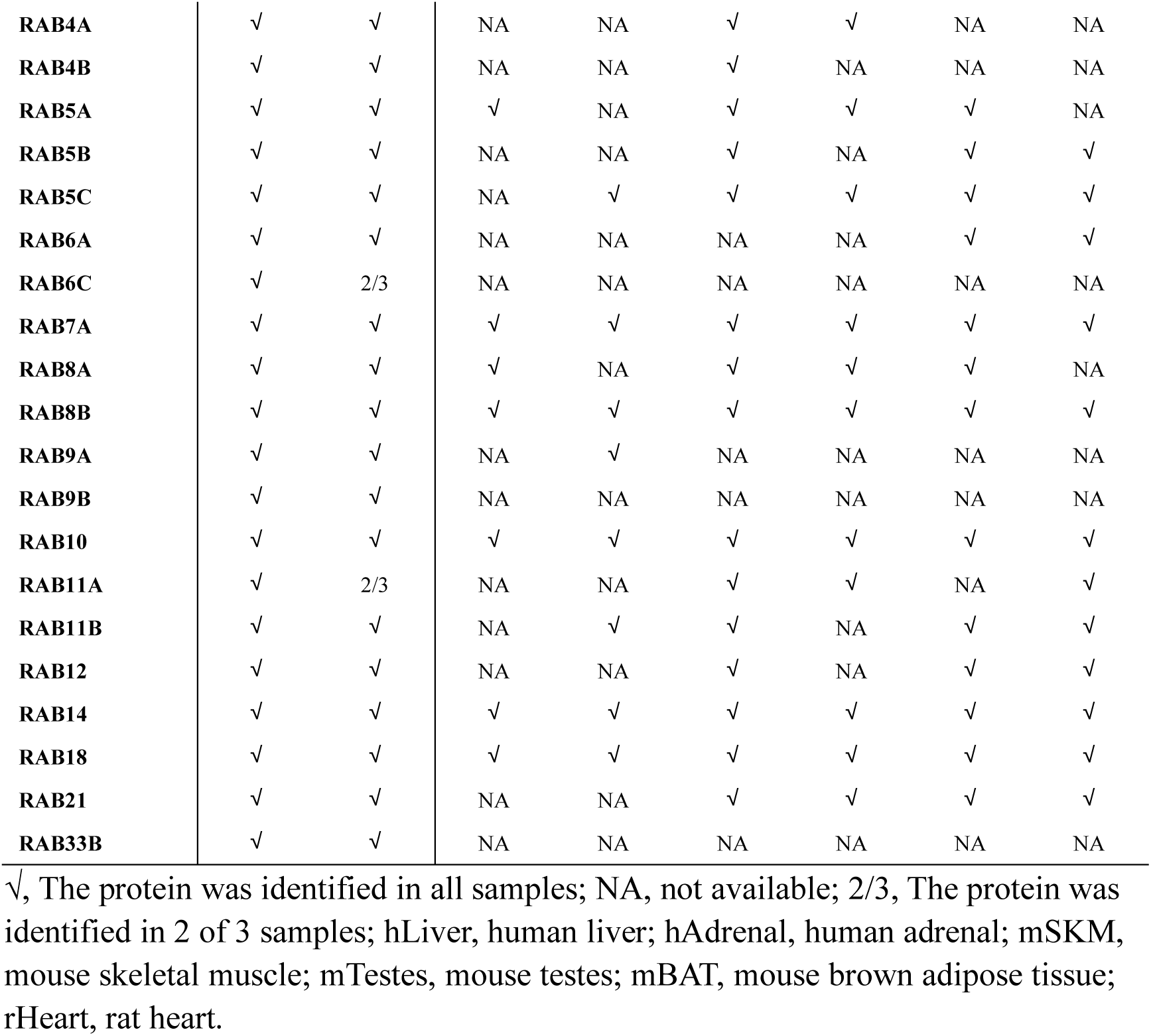
Major LD-associated proteins in skeletal muscle identified by MS.

Previous studies from our laboratory and others have demonstrated that in oxidative tissues^38^, such as mouse skeletal muscle, heart, and BAT, LDs and mitochondria interact closely, potentially through anchoring mechanisms. To investigate whether this occurs in human skeletal muscle, we examined mitochondrial subcompartment proteins identified in the LD proteome (Table 3). Consistently, our analysis revealed that LDs isolated from human skeletal muscle also contain numerous mitochondrial proteins, including components from the outer membrane, inner membrane, and matrix. Most of these proteins were also detected in the LD proteomes of mouse skeletal muscle^43^, mouse BAT^39^, and rat heart^44^. These findings suggest that LDAM are present in human skeletal muscle, with mitochondria maintaining structural integrity and functionality within this association.

**Table 3.**
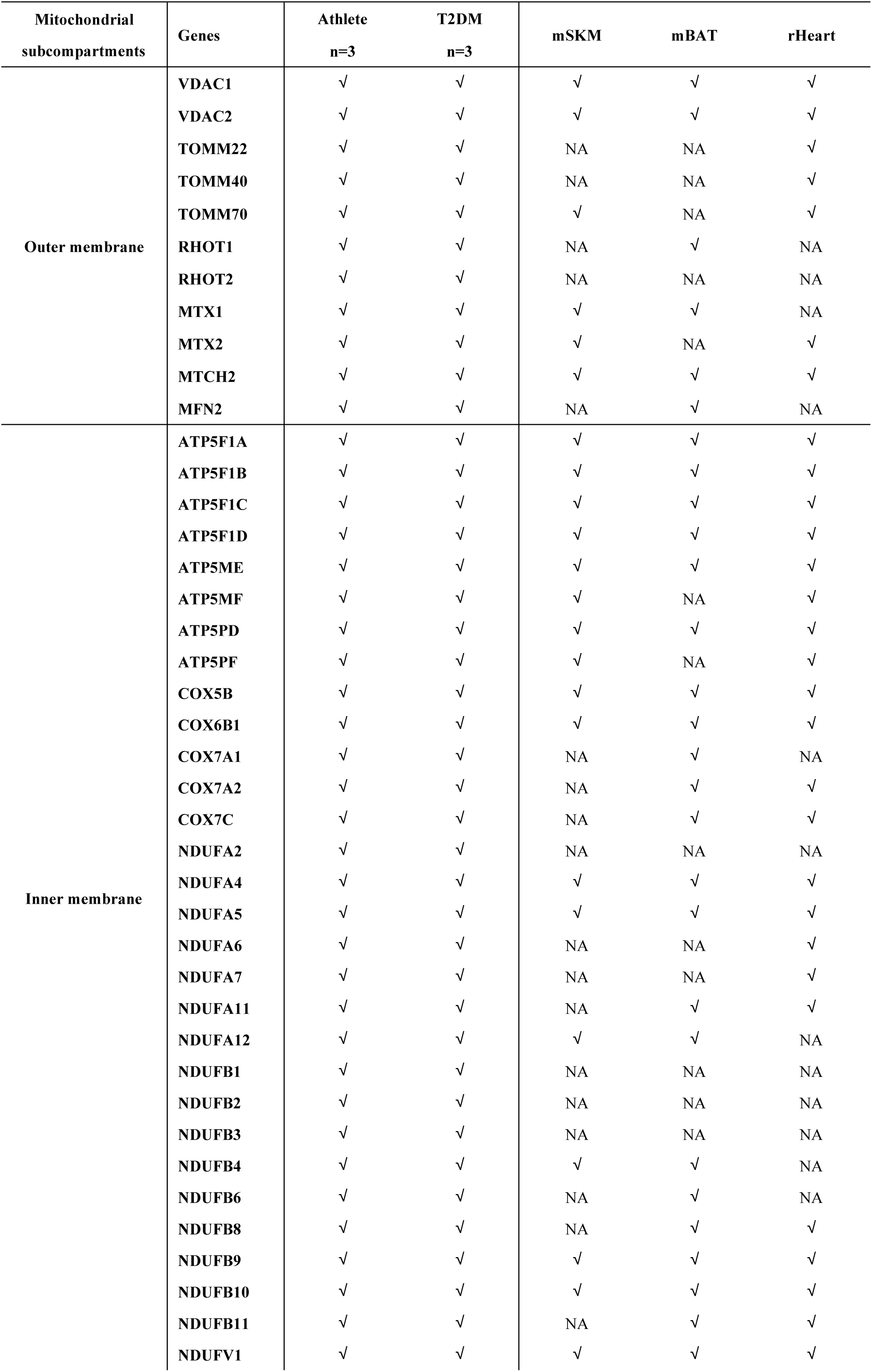
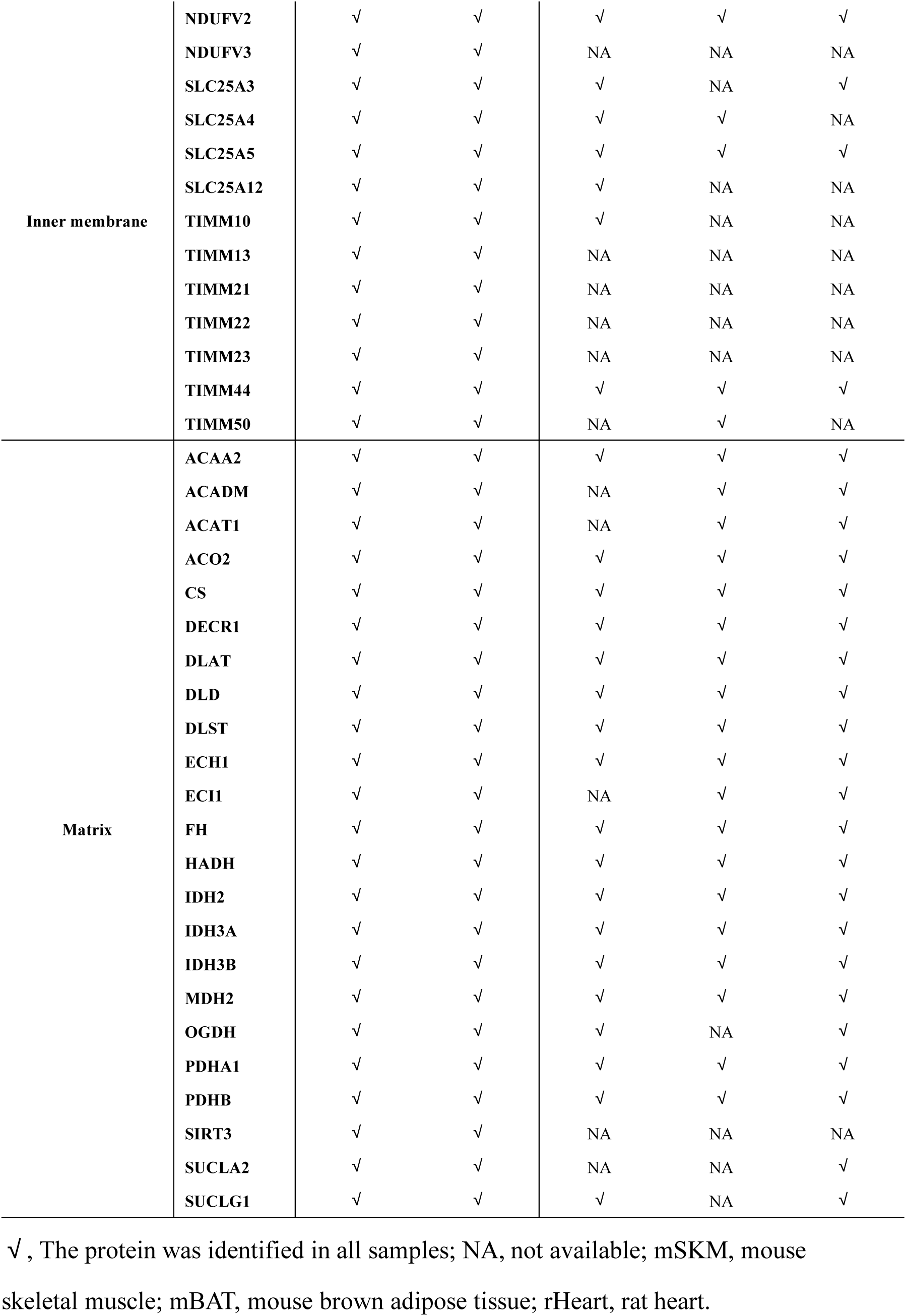
Major mitochondrial subcompartment proteins on LDs identified by MS.

### Comparative proteomic analysis of LDs and tissues from human skeletal muscle

Quantification of skeletal muscle TAG content showed that intramyocellular lipid (IMCL) levels were similar between endurance-trained athletes and T2DM patients, consistent with the athlete’s paradox (Fig. S1D). Additionally, most LDs exhibited diameters ranging from 0.1 to 3 µm, with no significant differences in size distribution between skeletal muscle samples from athletes and T2DM patients (Fig. 1Bb, c). The skeletal muscle LDs were rich in TAG, and the relative amounts of major lipid classes showed minimal differences between the two groups (Fig. 1Bd). These findings underscore the importance of investigating differences in LD properties between athletes and T2DM patients to better understand insulin signaling in human skeletal muscle. To this end, we performed quantitative proteomic analysis using data-independent acquisition (DIA) on LDs isolated from skeletal muscles of athletes and T2DM patients. The proteins extracted from skeletal muscle LDs of athletes and T2DM patients (n=3 per group) were digested with trypsin, and subjected to Orbitrap LC-MS. Proteins extracted from skeletal muscle tissues were analyzed in parallel. Principal component analysis (PCA) and protein clustering analysis classified the three athlete LD samples into one cluster, and similarly classified the three T2DM patient LD samples into another distinct cluster (Fig. 2A). The tissue samples exhibited an analogous clustering pattern (Fig. 2B).

**Figure 2.**
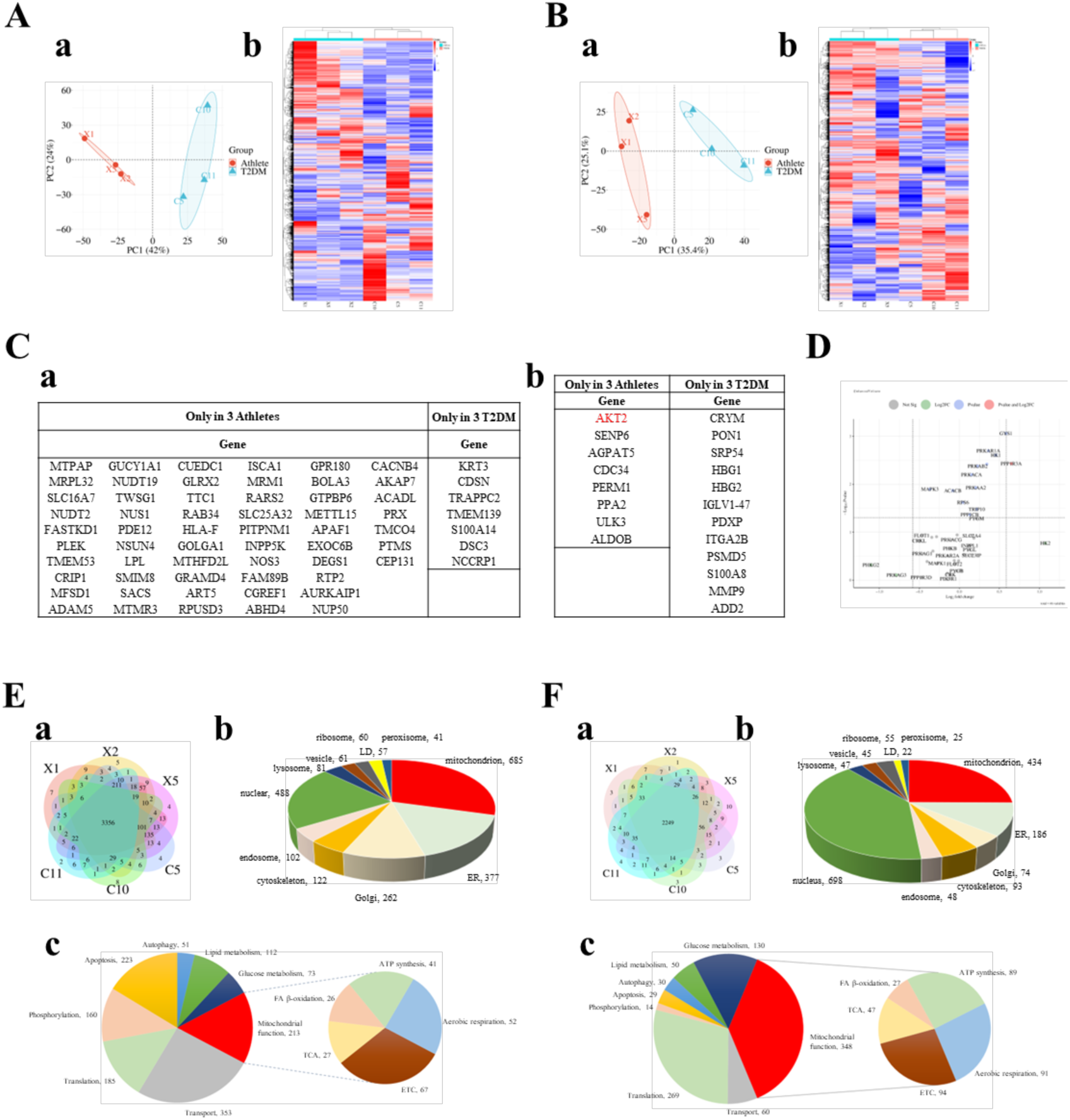
Comparative proteomic analysis of LDs and tissues from human skeletal muscle. (A) PCA factor map (a) and clustering analysis (b) of skeletal muscle LD proteome from 3 athletes and 3 T2DM patients. (B) PCA factor map (a) and clustering analysis (b) of skeletal muscle proteome from 3 athletes and 3 T2DM patients. (C) The proteins only present in 3 athletes or only in 3 T2DM patients. LD (a) and tissue (b) proteomes are shown in the table. (D) Volcano plot of insulin signaling pathway proteins in tissue proteome. (E) Venn diagram shows the overlap and distribution of identified LD proteins between athletes and T2DM patients skeletal muscle samples (a). Proteins with reproducible identification (present in all six LD samples) and containing more than 3 unique peptides were selected for cellular component analysis (b) and functional clustering analysis (c). GOCC, GOBP and KEGG databases were used for initial clustering. (F) Venn diagram shows the overlap and distribution of identified tissue proteins between athletes and T2DM patients skeletal muscle samples (a). Proteins with reproducible identification (present in all six tissue samples) and containing more than 3 unique peptides were selected for cellular component analysis (b) and functional clustering analysis (c). GOCC, GOBP and KEGG databases were used for initial clustering.

Mass spectrometry identified 4,030, 4,034, and 4,068 proteins in the three athlete LD samples (Fig. S2Aa), and 3,984, 3,623, and 3,739 proteins in the three T2DM patient LD samples (Fig. S2Ab). Among these, 3,907 proteins (approximately 94% of total identified proteins) were consistently detected across all athlete replicates, while 3,436 proteins (approximately 84% of total identifications) were consistently detected in T2DM patient replicates. Proteins exclusively identified in either athlete or T2DM patient LDs are listed in the table (Fig. 2Ca). GO cellular component (GOCC) analysis revealed that proteins exclusively identified in athlete skeletal muscle LDs were primarily enriched in mitochondria (Fig. S2C).

In the skeletal muscle tissue proteome, 2,559, 2,509, and 2,541 proteins were identified in the three athlete samples, with 2,399 proteins (91%) reproducibly detected as shown in the Venn diagram (Fig. S2Ba). Correspondingly, 2,516, 2,524, and 2,482 proteins were identified in the three T2DM tissue samples, with 2,361 proteins (90%) reproducibly detected (Fig. S2Bb). Proteins exclusively identified in either athlete or T2DM patient tissues are listed in the table (Fig. 2Cb). Among these, AKT2 was detected only in the skeletal muscle of athletes (Fig. 2Cb). In addition, quantitative analysis revealed that multiple proteins involved in insulin signaling pathways were upregulated in athlete skeletal muscle tissues, while only two proteins were slightly downregulated (Fig. 2D). The upregulation of insulin signaling pathway proteins suggests that athletes exhibit higher insulin sensitivity in skeletal muscle compared to T2DM patients.

Collectively, these results demonstrate high reproducibility across biological replicates and reliability of the mass spectrometry-based proteomic analysis, indicating the accuracy of the LD purification and proteomics techniques.

Further analysis showed that 3,356 proteins (81%) were reproducibly identified in all six LD samples (Fig. 2Ea) and 2,249 proteins (85%) were reproducibly identified in all six tissue samples (Fig. 2Fa). To investigate the compositional characteristics and differential expression profiles between the LD proteome and tissue proteome in human skeletal muscle, we selected proteins that identified across all six samples with unique peptides ≥ 3 from the proteomic dataset for subsequent analysis. A total of 2,934 proteins were selected from the LD proteome and 1,792 proteins were selected from the tissue proteome. The selected proteins were clustered into pathways using the Kyoto Encyclopedia of Genes and Genomes (KEGG) pathway database, while their cellular components and biological process were functionally annotated based on the Gene Ontology (GO) database. KEGG pathway enrichment analysis revealed that the LD proteome and tissue proteome exhibited similar but subtly distinct enrichment profiles, with metabolic pathways being the most significantly enriched category in both datasets (Fig. S2D). GOCC analysis revealed that the LD proteome is predominantly enriched in mitochondria, ER, and Golgi apparatus (Fig. 2Eb), while the tissue proteome is primarily enriched in nucleus, followed by mitochondria and ER (Fig. 2Fb). The LD proteome contains numerous proteins derived from other organelles, such as mitochondria and ER, indicating that LDs in skeletal muscle exhibit extensive interactions with these organelles and suggesting active lipid metabolism in this tissue. GO biological process (GOBP) analysis revealed that the LD proteome is significantly enriched in processes related to mitochondrial energy metabolism, transport, and lipid metabolism (Fig. 2Ec). In contrast, the tissue proteome is predominantly associated with mitochondrial energy metabolism, translation, and glucose metabolism (Fig. 2Fc). These findings reveal that the LD proteome and the tissue proteome exhibit notable differences in cellular components and biological functions.

### Comparative proteomic analysis of skeletal muscle LDs from athletes and T2DM patients

We quantified the protein abundance ratio in the proteomes between the athlete group and the T2DM patient group, followed by statistical significance analysis of the observed differences. The ratio value was defined as **Ratio = Protein expression in athletes/Protein expression in T2DM patients**. The calculated *p* value indicates whether the observed protein abundance ratio (Athletes/T2DM patients) reaches statistical significance, with *p* < 0.05 considered statistically significant. In the LD proteome, among the 2,934 proteins identified across all six samples with at least three unique peptides, 1,984 proteins showed statistically significant differences (*p* < 0.05). In the tissue proteome, out of 1,792 proteins, 805 showed significant differences (*p* < 0.05).

Next, GOCC annotation and KEGG pathway enrichment analysis of significantly differentially expressed proteins revealed distinct patterns between LD and tissue proteomes. In the LD proteome, most of mitochondrial proteins, ER resident proteins, and LD-associated proteins were significantly upregulated in athletes, whereas cytoskeletal proteins were markedly downregulated (Fig. 3A). Proteins associated with key metabolic pathways, including fatty acid metabolism, glycerolipid metabolism, glycolysis/gluconeogenesis, insulin signaling pathway, and glucagon signaling pathway, exhibited predominant upregulation (Fig. 3B). By comparison, the tissue proteome showed consistent upregulation of mitochondrial and ER proteins, but exhibited mixed alterations (both increases and decreases) in LD-associated proteins and cytoskeletal proteins (Fig. S3A). Notably, proteins associated with the same metabolic pathways in the LD proteome also showed consistent upregulation (Fig. S3B). These findings indicate that, compared to T2DM patients, athletes exhibit enhanced lipid metabolism activity in both skeletal muscle tissue and LDs.

**Figure 3.**
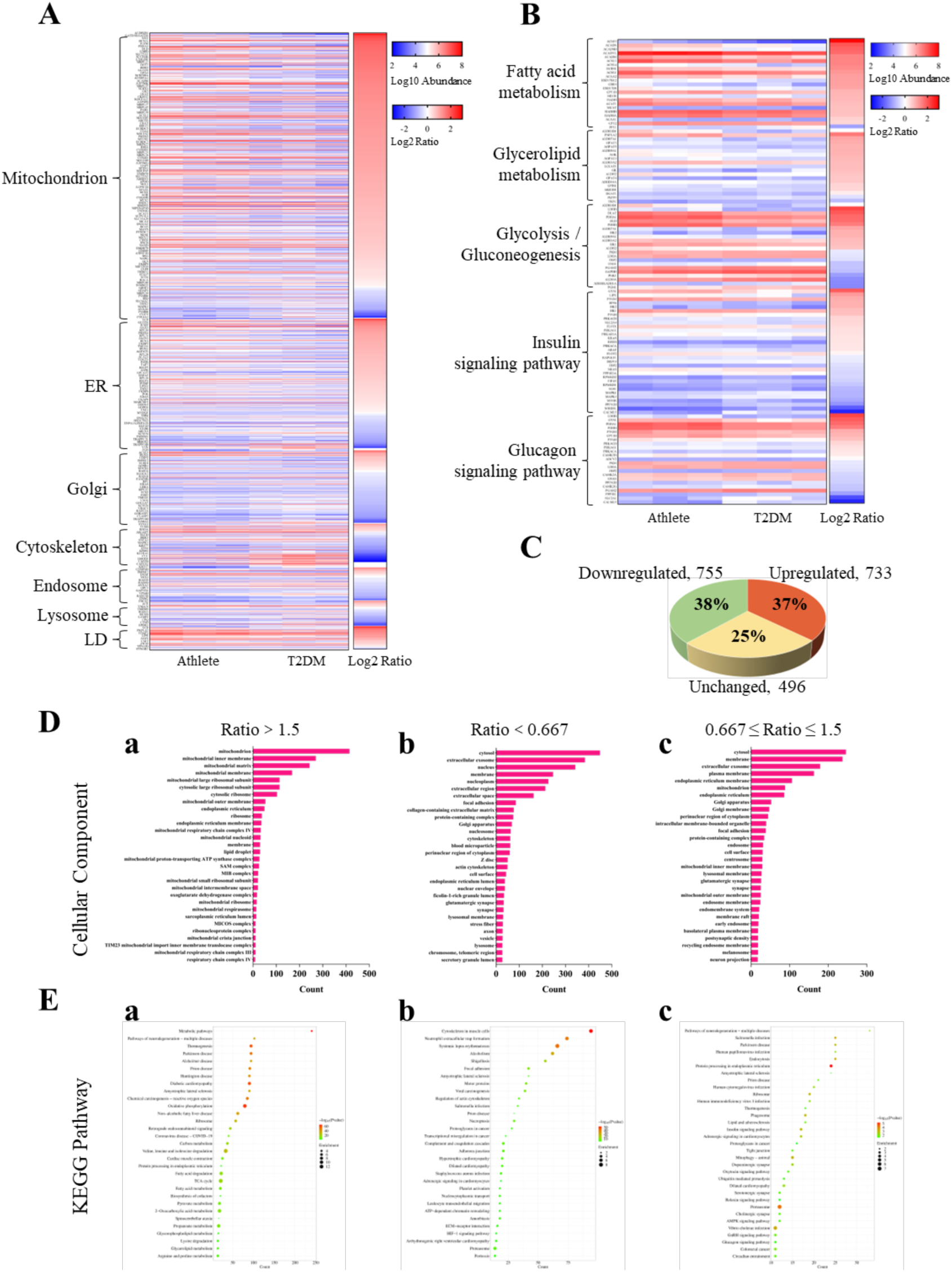
Comparative proteomic analysis of skeletal muscle LDs from athletes and T2DM patients. (A) The heatmap of differentially expressed proteins in the LD proteome. Proteins with unique peptide ≥ 3 and *p* < 0.05 were selected for analysis by GOCC enrichment. The color gradient represents the log10-transformed abundance of each protein, while the ratio color intensity reflects the log2-transformed fold change (athletes vs. T2DM patients). The heatmap of differentially expressed functional pathway proteins in the LD proteome. Proteins with unique peptide ≥ 3 and *p* < 0.05 were initially selected by KEGG pathway enrichment and subsequently filtered to specifically identify those involved in fatty acid metabolism, glycerolipid metabolism, glycolysis/gluconeogenesis, insulin signaling pathway, and glucagon signaling pathway for further analysis. (B) Percentages of upregulated, downregulated, and unchanged proteins relative to the total protein (*p* < 0.05) in the LD proteome. (C) Top 30 GOCC enrichment categories for upregulated (Ratio > 1.5, a), downregulated (Ratio < 0.667, b), and unchanged (0.667 ≤ Ratio ≤ 1.5, c) proteins identified in the LD proteome. (D) Top 30 KEGG enrichment pathways for upregulated (Ratio > 1.5, a), downregulated (Ratio < 0.667, b), and unchanged (0.667 ≤ Ratio ≤ 1.5, c) proteins identified in the LD proteome.

To further characterize the differentially expressed proteins in the LD and tissue proteomes between athletes and T2DM patients, we defined proteins with a Ratio > 1.5 as upregulated in athletes, those with 0.667 ≤ Ratio ≤ 1.5 as unchanged, and those with a Ratio < 0.667 as downregulated. In the LD proteome, 733 proteins (37%) were upregulated, 755 proteins (38%) were downregulated, and 496 proteins (25%) remained unchanged (Fig. 3C). In the tissue proteome, 303 proteins (38%) were upregulated, 147 proteins (18%) were downregulated, and 355 proteins (44%) showed no change (Fig. S3C). These proteins were subsequently subjected to GOCC annotation and KEGG pathway enrichment analysis. In the LD proteome, GOCC analysis showed that upregulated proteins were primarily localized to mitochondria, while downregulated proteins were predominantly located in the cytosol (Fig. 3D). Among the top 50 upregulated proteins, the majority were mitochondrial proteins (Table S1), while the top 50 downregulated proteins included many cytoskeletal proteins (Table S2). KEGG pathway analysis showed that upregulated proteins were enriched in metabolic pathways, whereas downregulated proteins were enriched in the cytoskeleton in muscle cells (Fig. 3E). By comparison, in the tissue proteome, upregulated proteins were similarly localized to mitochondria and enriched in metabolic pathways (Fig. S3D, E).

However, downregulated proteins were predominantly localized to extracellular exosomes and the extracellular region, with pathway analysis showing enrichment in systemic lupus erythematosus and neutrophil extracellular trap formation (Fig. S3D, E). These results indicate that, compared to T2DM patients, athletes exhibit similar profiles of upregulated proteins in both LDs and skeletal muscle tissue, but distinct profiles of downregulated proteins. The increase in mitochondrial proteins within the LD proteome suggests that skeletal muscle LDs in athletes exhibit enhanced lipid metabolism activity and greater energy utilization efficiency.

### Enhanced activity of LD-anchored mitochondria in skeletal muscle from athletes compared to T2DM patients

Compared to T2DM patients, an increased abundance of mitochondrial proteins was detected in the LD proteome of athlete skeletal muscle. However, it remains unclear whether this elevation in mitochondrial proteins is attributed to enhanced LD-mitochondria contacts or to increased activity of LDAM in skeletal muscle of athletes. To elucidate the underlying mechanisms responsible for the observed increase in mitochondrial proteins, we conducted comprehensive analyses of the detected mitochondrial proteins based on their intra-mitochondria localization (including inner membrane, matrix, outer membrane, and intermembrane space) and functional classification (including electron transport and ATP synthesis, TCA cycle, fatty acid beta-oxidation, cristae formation, mitochondrial translation, and mitochondrial organization).

Analysis of the mitochondrial sub-compartments in the LD proteome revealed distinct distribution patterns of protein ratios across different mitochondrial sub-compartments (Fig. 4A). Most mitochondrial proteins were upregulated (Ratio > 1), but the percentage distribution of abundance Ratio for mitochondrial proteins were different across the four mitochondrial sub-compartments in the LD proteome. For mitochondrial inner membrane and matrix proteins, the proportions with Ratio > 2 were 59% and 67% respectively, with over 10% of proteins were Ratio > 3 (Fig. 4C). Mitochondrial outer membrane and intermembrane space proteins showed lower proportions at 27% and 39% for Ratio > 2, with only a minimal fraction reaching Ratio > 3 (Fig. 4C). In contrast, while most mitochondrial proteins were also upregulated, their Ratio primarily ranged between 1-2 in the tissue proteome (Fig. S4A, C). Specifically, only 33% and 26% of inner membrane and matrix proteins exceeded Ratio > 2, approximately half the proportion observed in the LD proteome, and less than 5% achieving Ratio > 3 (Fig. S4C). For outer membrane and intermembrane space proteins, the proportions with Ratio > 2 were 22% and 13% (Fig. S4C). Notably, the LD proteome showed significantly higher proportions of inner membrane and matrix proteins with Ratio > 2 compared to their counterparts in the tissue proteome. A higher ratio of mitochondrial inner membrane and matrix proteins suggests potential enhancement of LDAM activity. In addition, most mitochondrial proteins exhibited Ratio greater than 1 in the tissue proteome, indicating increased contacts between LDs and mitochondria in the skeletal muscle of athletes compared to T2DM patients.

**Figure 4.**
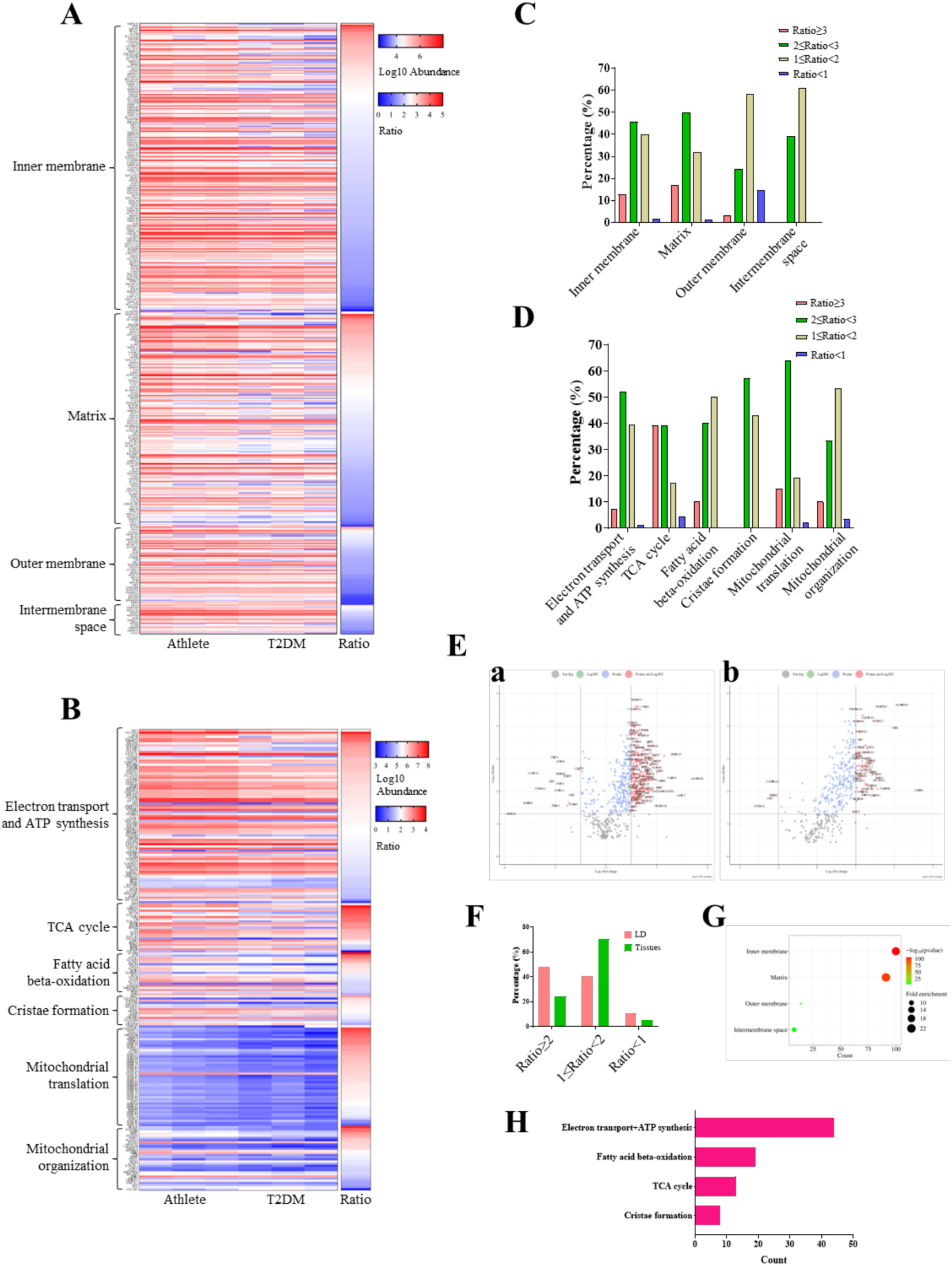
Enhanced activity of LDAM in athletes compared to T2DM patients. (A) The heatmap of differentially expressed mitochondrial subcompartment proteins in the LD proteome. Mitochondrial proteins with unique peptide ≥ 3 and *p* < 0.05 were selected for analysis by GOCC enrichment. The color gradient represents the log10-transformed abundance of each protein, while the ratio indicates the relative protein expression level in athletes compared to T2DM patients. (B) The heatmap of differentially expressed mitochondrial functional proteins in the LD proteome. Mitochondrial proteins with unique peptide ≥ 3 and *p* < 0.05 were initially selected by GOCC enrichment and subsequently filtered through GOBP enrichment to specifically identify those involved in electron transport and ATP synthesis, TCA cycle, fatty acid beta-oxidation, cristae formation, mitochondrial translation, and mitochondrial organization for further analysis. (C) The percentage distribution of abundance ratios for mitochondrial proteins across the four mitochondrial subcompartments in the LD proteome. The data is derived from the results in Figure A. (D) The percentage distribution of abundance ratios for mitochondrial functional proteins in the LD proteome. The data is derived from the results in Figure B. (E) Volcano plot of mitochondrial proteins in the LD proteome (a) or tissue proteome (b) from human skeletal muscle. Mitochondrial proteins were selected for analysis by GOCC enrichment. Proteins with ratio ≥ 2 or ratio ≤ 0.5 were annotated in red. Thresholds: *p* value < 0.05. (F) The percentage distribution of mitochondrial proteins across different abundance ratios in both LD and whole-tissue proteomes. (G) Subcompartmental distribution of mitochondrial proteins with a ratio ≥ 2 in the LD proteome but not in the tissue proteome. GOCC analysis was employed to determine the subcompartmental localization of these proteins. (H) Functional enrichment analysis of mitochondrial proteins with a ratio ≥ 2 in the LD proteome but not in the tissue proteome by GOBP analysis.

From a functional perspective, the LD proteome analysis revealed that proteins associated with key mitochondrial processes, including electron transport and ATP synthesis, TCA cycle, fatty acid beta-oxidation, cristae formation, mitochondrial translation, and mitochondrial organization, predominantly exhibited Ratio greater than 1 (Fig. 4B). This observation indicates enhanced functional activity of LDAM in athletes. It is worth noting that, among these functional categories, the proportion of TCA cycle related proteins with Ratio > 3 reached as high as 39%, and the proportion of fatty acid beta-oxidation related proteins with Ratio > 3 was 10% (Fig. 4D). Beyond mitochondrial organization, over 50% of proteins in other mitochondrial functional categories exhibited Ratio > 2 (Fig. 4D). In contrast, mitochondrial functional proteins in the tissue proteome primarily displayed ratio ranging from 1 to 2, with only a very small proportion of proteins related to electron transport and ATP synthesis showing Ratio > 3 (Fig. S4B, D). Compared to the tissue proteome, the LD proteome exhibited significantly higher ratio of mitochondrial functional proteins. These findings suggest that, in contrast to T2DM patients, athletes display enhanced functional activity in LDAM, particularly increased electron transport and ATP synthesis and TCA cycle capacity.

To visualize the changes in mitochondrial proteins, we generated volcano plots of mitochondrial proteins in the LD proteome (Fig. 4Ea) and the tissue proteome (Fig. 4Eb), with mitochondrial proteins exhibiting Ratio > 2 or < 0.5 and *p* < 0.05 specifically annotated. Subsequent analysis revealed that 48% of mitochondrial proteins exhibited Ratio > 2 in the LD proteome, and only 24% in the tissue proteome (Fig. 4F). We further selected 204 proteins that exhibited Ratio > 2 specifically in the LD proteome, with no corresponding upregulation observed in the tissue proteome. These proteins showed more pronounced increases in mitochondria associated with LDs compared to those in the tissue, suggesting their critical role in distinguishing LDAM from cytosol mitochondria. GOCC analysis of these proteins revealed their predominant localization to the mitochondrial inner membrane and matrix (Fig. 4G), GOBP analysis showed significant enrichment in functions related to electron transport and ATP synthesis, fatty acid beta-oxidation, TCA cycle, and cristae formation (Fig. 4H). These findings further suggest that, in the skeletal muscle of athletes, LDAM exhibit enhanced lipid metabolic activity and improved energy utilization efficiency, highlighting their superior functional capacity in lipid handling and energy production.

### Reduced abundance of cytoskeletal proteins in the skeletal muscle LD proteome of athletes

GOCC analysis of the LD proteome revealed that most of microfilament and intermediate filament proteins were downregulated, while microtubule proteins exhibited partial downregulation (Fig. 5A). Specifically, 95% of microfilament proteins and 94% of intermediate filament proteins displayed Ratio < 1, whereas only 54% of microtubule proteins displayed Ratio < 1 (Fig. 5C). In contrast, tissue proteome analysis revealed that most microtubule, microfilament, and intermediate filament proteins were upregulated in athletes relative to T2DM patients (Fig. 5B, D). To systematically evaluate cytoskeletal protein regulation, we constructed volcano plots comparing expression ratio in the LD and tissue proteomes of athletes versus T2DM patients (Fig. 5E). Proteins meeting statistical significance (*p* < 0.05) were highlighted, with threshold lines demarcating Ratio > 1.5 (upregulated) or < 0.667 (downregulated). The analysis showed a pronounced downregulation of cytoskeletal proteins in the LD proteome of athletes, with the most of proteins clustering below the threshold line (Fig. 5Ea). In contrast, only a small subset of cytoskeletal proteins in the tissue proteome exhibited downregulation (Fig. 5Eb). We further selected proteins that exhibited Ratio < 0.5 specifically in the LD proteome, with no corresponding downregulation observed in the tissue proteome (Fig. 5F). GOCC analysis of these proteins revealed their predominant localization to the actin cytoskeleton (Fig. 5G). These findings indicate a compartment-specific decrease of cytoskeletal proteins on LDs in the skeletal muscle of athletes compared to T2DM patients, with no analogous reduction observed in the broader tissue proteome. This compartmentalized downregulation strongly implicates cytoskeletal proteins as critical regulators of LD function, potentially influencing lipid mobilization, LD morphology, or interorganellar interactions. The increased presence of cytoskeletal proteins on LDs in T2DM patient skeletal muscle may be closely associated with impaired LD migratory capacity.

**Figure 5.**
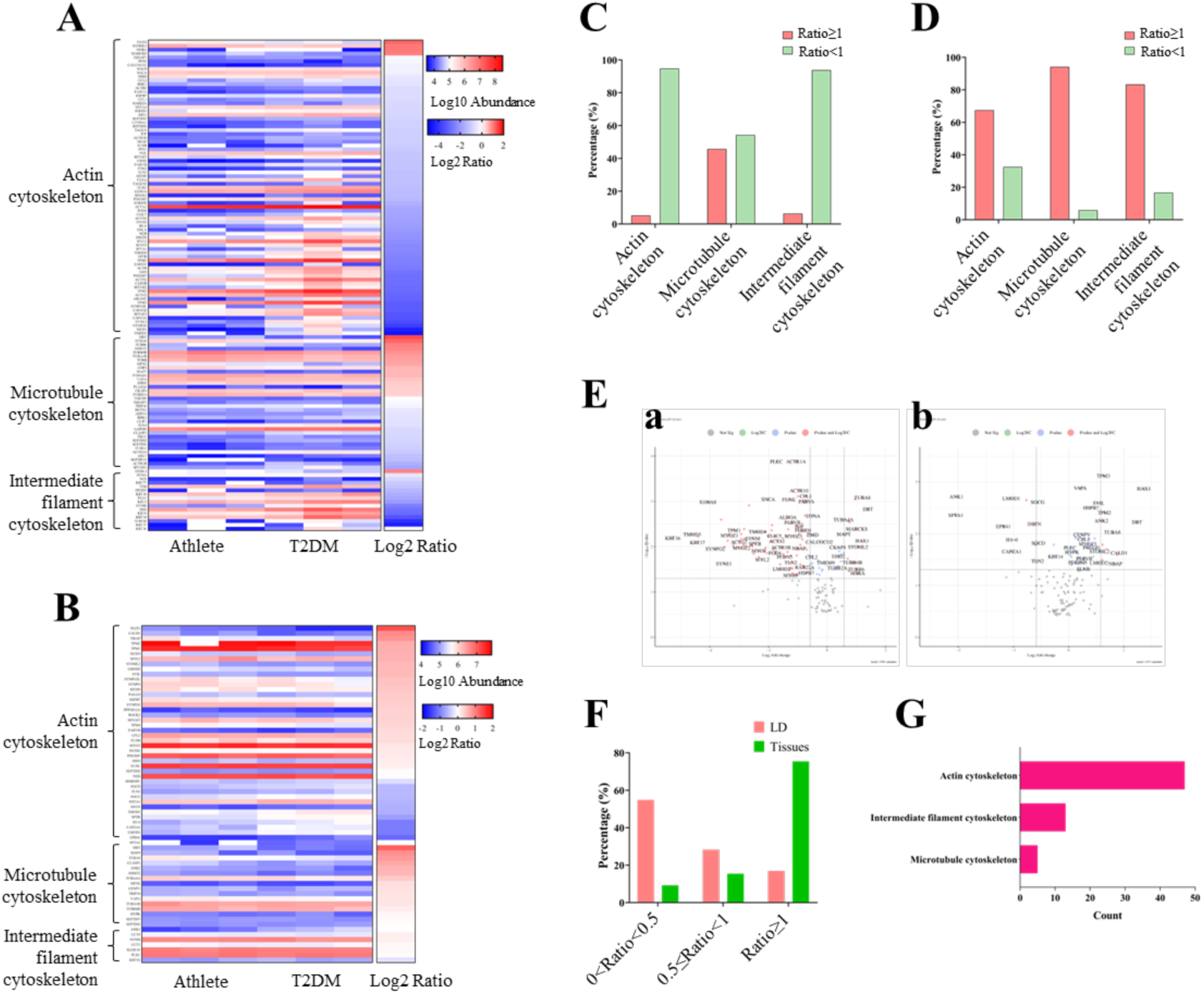
Reduced cytoskeletal proteins in the LD proteome of athlete skeletal muscle. (A) The heatmap of differentially expressed cytoskeletal proteins in the LD proteome. Cytoskeletal proteins with unique peptide ≥ 3 and *p* < 0.05 were selected for analysis by GOCC enrichment. The color gradient represents the log10-transformed abundance of each protein, while the ratio color intensity reflects the log2-transformed fold change (athletes vs. T2DM patients). (B) The heatmap of differentially expressed cytoskeletal proteins in the tissue proteome. Cytoskeletal proteins with unique peptide ≥ 3 and *p* < 0.05 were selected for analysis by GOCC enrichment. (C) The percentage distribution of abundance ratios for cytoskeletal proteins in the LD proteome. The data is derived from the results in Figure A. (D) The percentage distribution of abundance ratios for cytoskeletal proteins in the tissue proteome. The data is derived from the results in Figure B. (E) Volcano plot of cytoskeletal proteins in the LD proteome (a) or tissue proteome (b) from human skeletal muscle. Cytoskeletal proteins were selected for analysis by GOCC enrichment. Proteins with ratio > 1.5 or ratio < 0.667 were annotated in red. Thresholds: *p* value < 0.05. (F) The percentage distribution of cytoskeletal proteins across different abundance ratios in both LD and whole-tissue proteomes. (G) The compartmental distribution of cytoskeletal proteins with a ratio < 0.5 in the LD proteome but not in the tissue proteome.

### Up-regulation of ATGL in skeletal muscle LDs from athletes

The volcano plot analysis encompassed 2,934 proteins with unique peptide ≥ 3 that identified across all six LD samples, with LD-associated proteins were annotated and threshold lines demarcating Ratio > 1.5 (upregulated) or < 0.667 (downregulated) (Fig. 6A). Most of LD-associated proteins are upregulated in athletes (Table 4). The STRING web database was employed to analyze the physical and functional interactions of all the identified upregulated LD-associated proteins. The interaction network of upregulated LD-associated proteins showed a predominant association with TAG mobilization in skeletal muscle (Fig. 6B). It is noteworthy that there is a 2.97-fold increase in ATGL (PNPLA2) in skeletal muscle LDs of athletes (Table 4).

**Figure 6.**
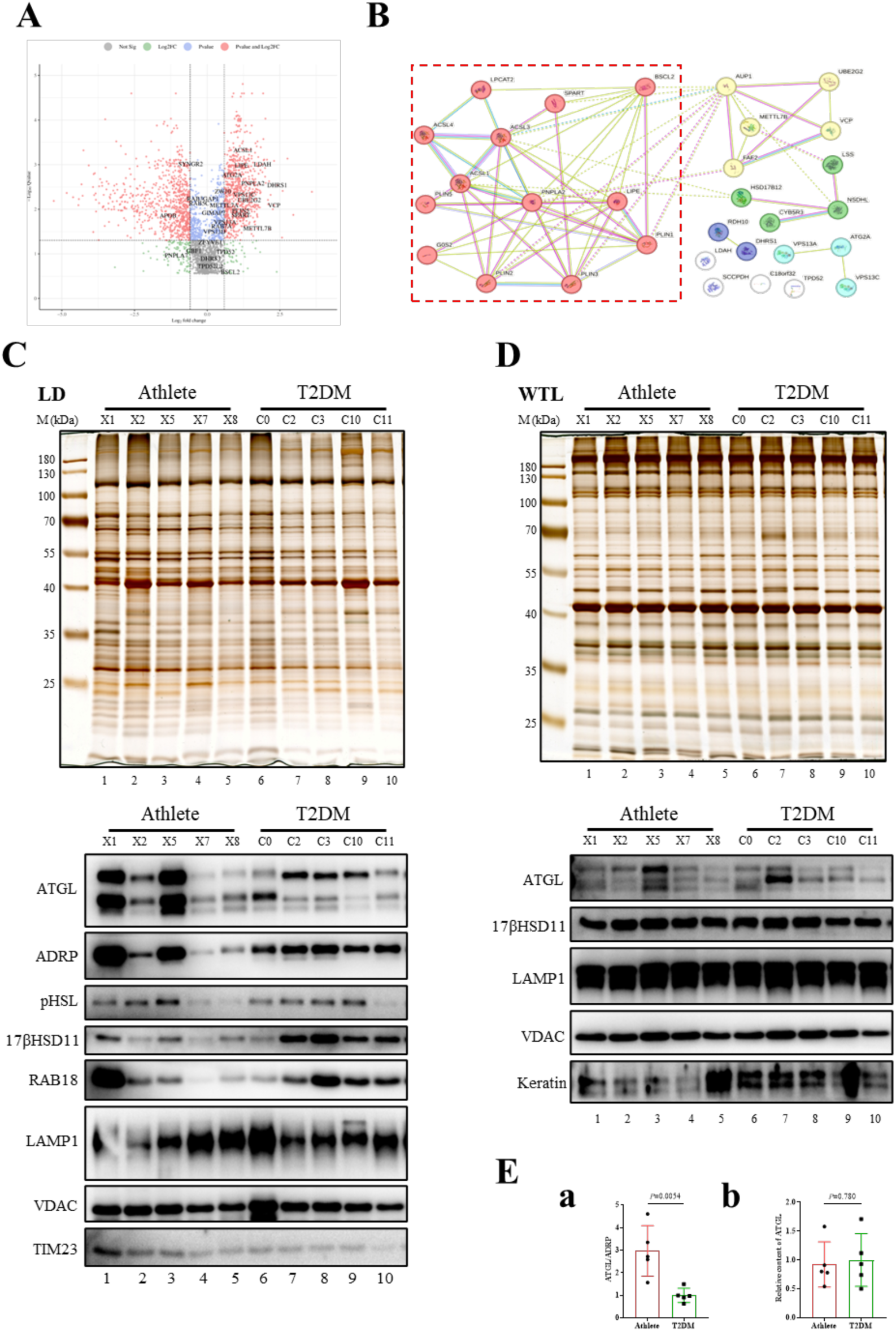
Up-regulation of ATGL in skeletal muscle LDs from athletes. (A) The differentially expressed proteins in the LD proteome analyzed by volcano plots. LD-associated proteins were annotated. (B) The interaction network of LD-associated proteins with upregulated expression in the LD proteome from skeletal muscle of athletes compared with T2DM patients using STRING interaction network. TAG mobilization related proteins such as HSL (LIPE), ATGL (PNPLA2), ACSL3, and ACSL4 were upregulated. (C) Silver staining of proteins was performed to verify consistent sample loading across all LD samples. LD-associated protein ATGL, ADRP, pHSL, 17βHSD11, and RAB18 levels from athletes and T2DM patients were validated by immunoblotting. Lysosomal protein LAMP1 and mitochondrial protein VDAC and TIM23 were validated by immunoblotting using the corresponding antibody. (D) Silver staining of proteins was performed to verify consistent sample loading across all tissue samples. LD-associated protein ATGL and 17βHSD11, lysosomal protein LAMP1 and mitochondrial protein VDAC were validated by immunoblotting using the corresponding antibody. (E) The relative protein level of ATGL in LDs (a) and tissues (b). Grayscale analysis was conducted utilizing Image J software.

**Table 4.** Differential LD-associated proteins in athletes and T2DM skeletal muscles <u>identified by quantitative MS.</u>.

| Genes | Protein Descriptions | Unique Peptides | Ratio | Q value |
| --- | --- | --- | --- | --- |
| DHRS1 | Dehydrogenase/reductase SDR family member 1 | 7 | 5.31 | 0.0029 |
| VCP | Transitional endoplasmic reticulum ATPase | 43 | 4.93 | 0.0082 |
| G0S2 | G0/G1 switch protein 2 | 6 | 4.37 | 0.0024 |
| LDAH | Lipid droplet-associated hydrolase | 6 | 3.74 | 0.0010 |
| METTL7B | Thiol S-methyltransferase METTL7B | 7 | 3.31 | 0.0268 |
| SCCPDH | Saccharopine dehydrogenase-like oxidoreductase | 32 | 3.12 | 0.0039 |
| **PNPLA2** | **Patatin-like phospholipase domain-containing protein 2** | **18** | **2.97** | **0.0026** |
| UBE2G2 | Ubiquitin-conjugating enzyme E2 G2 | 4 | 2.72 | 0.0062 |
| ACSL3 | Fatty acid CoA ligase Acsl3 | 46 | 2.70 | 0.0059 |
| ACSL4 | Long-chain-fatty-acid--CoA ligase 4 | 20 | 2.70 | 0.0011 |
| CYB5R3 | NADH-cytochrome b5 reductase 3 | 35 | 2.66 | 0.0101 |
| LDAF1 | Lipid droplet assembly factor 1 | 7 | 2.57 | 0.0062 |
| LSS | Lanosterol synthase | 30 | 2.48 | 0.0099 |
| PLIN2 | Perilipin-2 | 56 | 2.46 | 0.0115 |
| VPS13C | Intermembrane lipid transfer protein VPS13C | 194 | 2.42 | 0.0047 |
| ACSL1 | Long-chain-fatty-acid--CoA ligase 1 | 50 | 2.35 | 0.0005 |
| LIPE | Hormone-sensitive lipase | 22 | 2.26 | 0.0010 |
| NSDHL | Sterol-4-alpha-carboxylate 3-dehydrogenase, decarboxylating | 20 | 2.26 | 0.0131 |
| SPART | Spartin | 23 | 2.23 | 0.0135 |
| PLIN5 | Perilipin-5 | 35 | 2.22 | 0.0107 |
| HSD17B12 | Very-long-chain 3-oxoacyl-CoA reductase | 11 | 2.16 | 0.0008 |
| AUP1 | Lipid droplet-regulating VLDL assembly factor AUP1 | 20 | 2.14 | 0.0064 |
| PLIN3 | Perilipin-3 | 47 | 1.92 | 0.0104 |
| C18orf32 | UPF0729 protein C18orf32 | 3 | 1.87 | 0.0294 |
| PLIN1 | Perilipin-1 | 34 | 1.84 | 0.0127 |
| ATG2A | Autophagy-related protein 2 homolog A | 22 | 1.81 | 0.0017 |
| FAF2 | FAS-associated factor 2 | 10 | 1.63 | 0.0082 |
| LPCAT2 | Lysophosphatidylcholine acyltransferase 2 | 6 | 1.57 | 0.0087 |
| VPS13A | Intermembrane lipid transfer protein VPS13A | 112 | 1.53 | 0.0194 |
| RDH10 | Retinol dehydrogenase 10 | 4 | 1.50 | 0.0313 |
| METTL7A | Putative methyltransferase-like protein 7A | 19 | 1.50 | 0.0082 |
| ZW10 | Centromere/kinetochore protein zw10 homolog | 25 | 1.45 | 0.0040 |
| RAB7A | Ras-related protein Rab-7a | 15 | 1.41 | 0.0241 |
| PNPLA4 | Patatin-like phospholipase domain-containing protein 4 | 8 | 1.35 | 0.0171 |
| ABHD5 | 1-acylglycerol-3-phosphate O-acyltransferase ABHD5 | 15 | 1.28 | 0.0369 |
| BCAP31 | B-cell receptor-associated protein 31 | 17 | 1.23 | 0.0292 |
| VPS13D | Intermembrane lipid transfer protein VPS13D | 75 | 1.19 | 0.0316 |
| GIMAP7 | GTPase IMAP family member 7 | 8 | 1.17 | 0.0121 |
| RAB3GAP1 | Rab3 GTPase-activating protein catalytic subunit | 17 | 0.90 | 0.0058 |
| RAB5C | Ras-related protein Rab-5C | 8 | 0.83 | 0.0073 |
| HSD17B11 | Estradiol 17-beta-dehydrogenase 11 | 8 | 0.72 | 0.0070 |
| **SYNGR2** | Synaptogyrin-2 | 4 | 0.67 | 0.0010 |
| **APOB** | Apolipoprotein B-100 | 184 | 0.39 | 0.0138 |

To verify the proteomic results and validate the increase of ATGL on LDs, the LD fractions from five athletes and five T2DM patients were loaded with equal amounts of proteins and separated by SDS/PAGE (Fig. 6C). The results showed that ATGL content was significantly increased in the LD fractions of athletes compared to ADRP (Fig. 6C, Ea). Furthermore, we observed increased content of TIM23, a mitochondrial inner membrane protein, consistent with our proteomic findings (Fig. 6C). Notably, parallel analyses of whole tissue lysates showed no significant differences in total ATGL protein levels in skeletal muscle between athletes and T2DM patients (Fig. 6D, Eb). These findings indicate that the observed increase in ATGL specifically on LDs in athletes is attributable to enhanced subcellular localization of ATGL to LDs, rather than changes in total tissue ATGL protein levels.

### ATGL deficiency impairs AKT phosphorylation in C2C12 cells

To investigate the role of ATGL in insulin signaling within skeletal muscle cells, we generated ATGL knockout (KO) and overexpressing (OE) C2C12 cell lines (Figure 7A). Previous studies have shown that palmitic acid (PA) treatment impairs insulin signaling by reducing AKT phosphorylation^60,61^. Consistently, upon PA treatment followed by insulin stimulation, ATGL KO cells exhibited significantly reduced AKT phosphorylation compared to control cells, while total AKT levels remained unchanged (Figure 7A). In contrast, treatment with oleic acid (OA) did not affect AKT phosphorylation in WT cells, whereas in ATGL KO cells, OA treatment caused a slight decrease in p-AKT. Notably, in ATGL OE cells, treatment with either OA or PA did not alter p-AKT levels compared to controls (Figure 7A).

**Figure 7.**
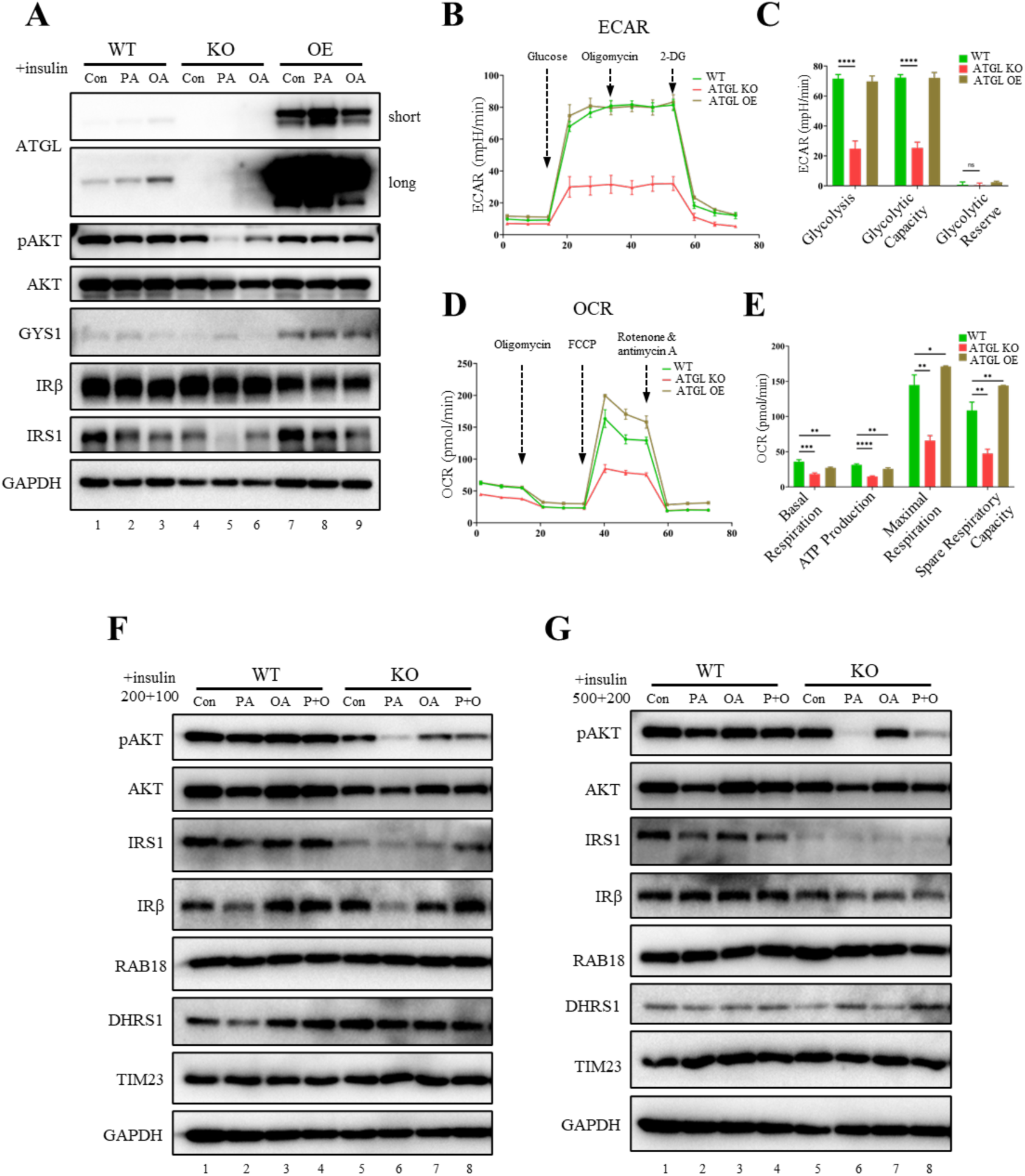
ATGL deficiency impairs AKT phosphorylation in myoblasts. (A) AKT phosphorylation levels were detected by immunoblotting in ATGL knockout and overexpression cells after PA or OA treatment. After treating the cells with PA for 20 hours, the culture medium was replaced with fresh medium containing 10 nM insulin, followed by an additional incubation for 15 minutes. (B) Cellular glycolytic activity was analyzed using the Seahorse XF96 Analyzer. ATGL knockout and overexpression C2C12 cells were seeded at an optimal density of 1×10^4^ cells in Seahorse culture plates. Glycolytic function was assessed by sequentially treating the cells with glucose (10 mM), oligomycin (1 μM), and 2-deoxyglucose (2-DG, 50 mM), while recording the extracellular acidification rate (ECAR) profiles. (C) Quantification of cellular glycolysis parameters in Seahorse assays. Glycolysis, glycolytic capacity, and glycolytic reserve were quantified in cells using Seahorse assay data. Data were analyzed by unpaired Student t-test and presented as mean ± SD. ****, *p* < 0.0001; ns, no significance. (D) Mitochondrial respiratory function was analyzed using the Seahorse XF96 Analyzer. ATGL knockout and overexpression cells were seeded at an optimal density of 1×10^4^ cells in Seahorse culture plates. Mitochondrial respiration was evaluated by measuring the oxygen consumption rate (OCR) under basal conditions and following sequential treatments with oligomycin (1.5 µM), the uncoupler FCCP (0.5 µM), and the Complex III and I inhibitors antimycin A and rotenone (0.5 µM each). (E) Quantification of mitochondrial function parameters in Seahorse assays. Basal respiration, ATP production, maximal respiration, and spare respiratory capacity were quantified from Seahorse assay data in cells. Data were analyzed by unpaired Student t-test and presented as mean ± SD. *, *p* < 0.05; **, *p* < 0.01; ***, *p* < 0.001. (F) AKT phosphorylation levels were detected by immunoblotting in ATGL knockout cells after PA, OA, or PA+OA treatment. PA was used at a concentration of 200 µM, OA at 100 µM, and the combination treatment (PA+OA) consisted of 200 µM PA and 100 µM OA. (F) AKT phosphorylation levels were detected by immunoblotting in ATGL knockout cells after PA, OA, or PA+OA treatment. PA was used at a concentration of 500 µM, OA at 200 µM, and the combination treatment (PA+OA) consisted of 500 µM PA and 200 µM OA.

Given the role of ATGL in mediating LD lipolysis, we next quantified cellular TAG levels under PA and OA treatment to explore whether lipid accumulation underlies p-AKT regulation. Both PA and OA increased TAG content (Figure S5A). ATGL KO led to further TAG accumulation, while ATGL OE reduced TAG levels, confirming effective modulation of intracellular lipid turnover. However, despite reduced TAG with ATGL OE, p-AKT did not increase further, suggesting additional regulatory factors in insulin signaling (Figure 7A). To clarify this, we examined ATGL distribution in OE cells and found that while total cellular ATGL was substantially increased, LD-associated ATGL increased only slightly (Figure S5B, C). This tight regulation of ATGL translocation to LDs may constrain the capacity to enhance p-AKT even when TAG levels decrease with ATGL OE.

To understand the mechanism underlying reduced p-AKT with ATGL deficiency, we examined upstream signaling components. ATGL KO cells displayed decreased IRS1 levels, likely contributing to the observed reduction in p-AKT (Figure 7A). Additionally, ATGL OE increased glycogen synthase 1 (GYS1) levels, indicating potential enhancement of downstream insulin signaling and glucose storage (Figure 7A).

We further assessed the metabolic state of these cells, as cellular energy status influences insulin responsiveness. ATGL KO cells exhibited reduced extracellular acidification rates (ECAR) and oxygen consumption rates (OCR), indicating decreased glycolytic capacity and impaired mitochondrial oxidative phosphorylation, respectively (Figure 7B-E). These metabolic impairments may further contribute to compromised insulin signaling under lipid overload in the absence of ATGL. We next investigated how the timing of PA treatment affects AKT phosphorylation. Short-term PA exposure increased AKT phosphorylation in both WT and ATGL KO cells under insulin-stimulated conditions, consistent with previous reports (Figure S6C, D)^62^. In contrast, long-term PA treatment reduced AKT phosphorylation regardless of the presence or absence of insulin in the culture medium and was accompanied by a slight decrease in IRS1 levels after 18 hours of treatment (Figure S6A-D). Given that OA has been reported to alleviate PA-induced insulin resistance^60,62^, we tested whether this protective effect persists in the absence of ATGL to assess whether ATGL is required for OA-mediated protection under lipid overload. Under low PA+OA co-treatment, ATGL KO cells exhibited significantly higher p-AKT levels compared to PA treatment alone, indicating that OA can effectively rescue PA-induced insulin resistance even in the absence of ATGL (Figure 7F). However, under high PA+OA conditions, only a slight increase in p-AKT was observed in ATGL KO cells, suggesting that the protective effect of OA is substantially attenuated when ATGL is absent under high lipotoxic stress (Figure 7G).

Collectively, these findings suggest that ATGL deficiency impairs AKT phosphorylation in skeletal muscle cells by reducing IRS1 abundance and impairing metabolic activity, thereby compromising insulin signaling under conditions of lipid overload. Furthermore, while OA can partially mitigate PA-induced insulin resistance, its protective effect is limited in the absence of ATGL, particularly under high PA conditions, underscoring the role of ATGL in maintaining insulin sensitivity during lipid stress.

### ATGL reduction and mitochondrial protein increase in skeletal muscle LDs of db/db mice

Due to the difficulty in sampling human skeletal muscle tissue, it is hard to maintain consistency in terms of gender, age, and tissue site. To address this limitation, we validated our findings in diabetic mice. The db/db mice exhibited characteristic features of T2DM, showing elevated fasting blood glucose levels approximately 8-fold higher than wild-type (WT) controls, along with a 1.8-fold increase in body weight (Fig. 8A, B). We collected skeletal muscle tissue from all four limbs of three mice, carefully removed the adipose tissue, and subsequently isolated and purified their LDs. The skeletal muscle weight of db/db mice was significantly lower than that of WT mice, and its proportion to body weight is about one-third of that of WT mice (Fig. 8C, D). However, the TAG content in db/db skeletal muscle was approximately three times that of WT mice (Fig. 8E), which is consistent with the lipid changes observed in skeletal muscle of diabetic patients compared to healthy individuals^27^. The size of LDs in skeletal muscle of db/db mice showed slight increase compared to WT controls (Fig. 8F, G). Equal amounts of protein from the LD fraction and five other fractions were separated by SDS-PAGE for subsequent analysis (Fig. 8H). ATGL was significantly reduced on LDs of db/db mice, while no change was observed in whole tissue lysates (Fig. 8H). Furthermore, the content of TIM23 was also decreased on LDs (Fig. 8H). These findings are consistent with those observed in human skeletal muscle LDs in the present work.

**Figure 8.**
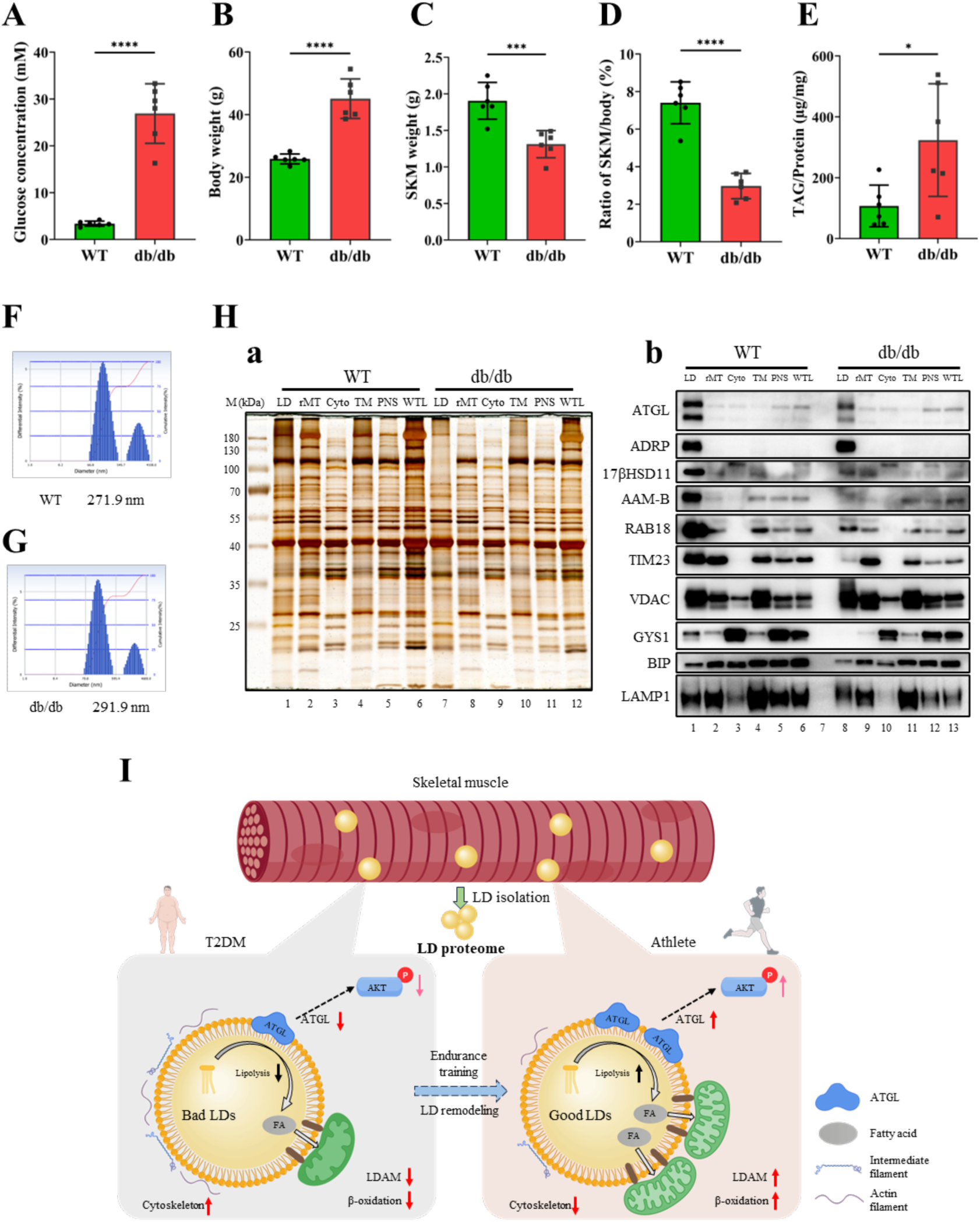
ATGL reduction and mitochondrial protein increase in skeletal muscle LDs of db/db mice. (A) The blood glucose levels in WT and db/db mice following 16-hour fasting. ****, *p* < 0.0001. (B) Body weight of WT and db/db mice. ****, *p* < 0.0001. (C) Skeletal muscle weight from four limbs in WT and db/db mice after removal of adipose tissue. ***, *p* < 0.001. (D) Ratio of skeletal muscle mass from four limbs to body weight in mice. ****, *p* < 0.0001. (E) TAG content of skeletal muscle tissues in WT and db/db mice. *, *p* < 0.05. (F) The size of LDs from skeletal muscle in WT mice. Skeletal muscles from four limbs of three WT mice were mixed for LD isolation. (G) The size of LDs from skeletal muscle in db/db mice. Skeletal muscles from four limbs of three db/db mice were mixed for LD isolation. (H) Protein fractions isolated from skeletal muscle, including lipid droplets (LDs), rough mitochondria (rMT), cytosol (Cyto), total membrane (TM), and whole tissue lysate (WTL), were subjected to silver staining (a) and further immunoblot analysis (b). Protein loading normalization based on equivalent TAG content for LD samples. ATGL expression along with other proteins of interest (ADRP, TIM23, BIP, and LAMP1) was assessed by Western blot analysis. (I) The schematic illustrates the differences in the protein profiles of “bad LDs” in T2DM patients and “good LDs” in athletes. The “bad LDs” in the skeletal muscle of T2DM patients are characterized by higher levels of cytoskeletal proteins, reduced LDAM, and lower ATGL compared to the “good LDs” observed in athletes. Additionally, LDAM in skeletal muscle of athletes exhibits enhanced energy metabolism activity, which is important for efficient utilization of fatty acids. Furthermore, elevated ATGL localization to LDs may promote AKT phosphorylation via an undefined mechanism, leading to enhanced insulin sensitivity in skeletal muscle. Image was prepared with BioGDP.com.

Next, we conducted parallel experiments in ob/ob mice, another well-established model of metabolic dysfunction. The ob/ob mice exhibited significantly higher body weight but lower skeletal muscle mass compared to WT controls, with skeletal muscle TAG content approximately 3-fold higher than that in WT mice (Fig. S7A-D). Although fasting blood glucose levels showed only a slight increase in ob/ob mice compared to WT controls, both glucose tolerance and insulin tolerance tests revealed significant impairment, indicating compromised insulin responsiveness (Fig. S7E, F). The skeletal muscle LDs of ob/ob mice were slightly larger in size than those of WT (Fig. S7G, H), and both ATGL and TIM23 protein levels on LDs were significantly decreased (Fig. S7I). These findings were consistent with the results observed in LD samples of db/db mice and human.

In the present study, we isolated LDs from human skeletal muscle, and established the proteome database of human skeletal muscle LD proteins from both athletes and T2DM patients. Comparative analysis of skeletal muscle LD proteins between athletes and T2DM patients revealed that athletes exhibited increased LDAM and significantly elevated ATGL levels on LDs, and showed reduced cytoskeletal proteins. Moreover, ATGL deficiency in myoblasts aggravates PA-induced insulin resistance, which suggests a potential regulatory role of LD-associated ATGL in the development of systemic insulin resistance (Fig 8I).

## Discussion

T2DM has emerged as a globally prevalent chronic disease that significantly impacts physical health and quality of life. As a metabolic disorder, skeletal muscle insulin resistance is widely recognized as a key contributor to T2DM pathogenesis. Lipid metabolism within skeletal muscle is closely linked to insulin sensitivity. It is now recognized that skeletal muscle insulin sensitivity is influenced not only by the total intramyocellular lipid (IMCL) content but also by the physiological state of LDs. However, the contribution of the proteomic composition of LDs in human skeletal muscle to insulin sensitivity has not yet been explored. In this study, we profiled the LD proteome isolated from the skeletal muscle of seven endurance-trained athletes and nine T2DM patients, establishing the first comprehensive database of human skeletal muscle LD-associated proteins in both athletes and T2DM patients. LD proteomic analysis revealed numerous proteins that are typically localized to other organelles, including mitochondria, ER, lysosomes, and Golgi apparatus. This finding strongly suggests extensive interactions between LDs and multiple cellular organelles in skeletal muscle. Previous studies have consistently demonstrated this phenomenon^43^, indicating that LDs represent highly active lipid metabolic hubs in skeletal muscle. We believe these findings will significantly advance our understanding of how lipid metabolism in human skeletal muscle contributes to insulin signaling.

Generally, IMCL is negatively correlated with insulin sensitivity, however, endurance-trained athletes have increased IMCL level and enhanced insulin sensitivity. This phenomenon, known as the athlete’s paradox, provides an excellent biological model for investigating the association between skeletal muscle LDs and insulin resistance. Therefore, we provide the first comparative analysis of the LD proteome of skeletal muscle between these two distinct metabolic groups. Comparative proteomic analysis identified 733 upregulated and 755 downregulated proteins in LDs of athletes. The upregulated proteins were primarily enriched in mitochondria while downregulated proteins comprised numerous cytoskeletal proteins. Our study provides direct evidence that the interactions between LD and mitochondria were enhanced in athlete skeletal muscle, and these LDAM exhibited greater oxidative capacity compared to T2DM. In support of this finding, morphological observation by electron microscope revealed that male endurance athletes have a greater LD-mitochondrion contact than untrained individuals^45^. This may partially explain the paradoxical coexistence of higher IMCL with improved insulin sensitivity in athletes. In addition, the cytoskeleton mediates intracellular LD trafficking, diminished LD-cytoskeleton interactions in athletes may reduce LD motility and inhibit LD fusion and growth, potentially enhancing lipid utilization efficiency within LDs.

Among many LD-associated proteins, the ATGL level in skeletal muscle LDs is 2.97-fold higher in athletes than in T2DM patients. Studies have reported that mice with global deletion of ATGL show increased skeletal muscle insulin sensitivity *in vivo*, while exhibiting reduced insulin sensitivity *in vitro*^57^, suggesting that the effect of ATGL deficiency on insulin sensitivity is much complex. In this study, ATGL was specifically upregulated in LDs of athletes, but not in total skeletal muscle tissues. ATGL deficiency aggravated PA-induced insulin resistance in C2C12 cells, whereas its overexpression failed to alleviate this effect, possibly due to unchanged ATGL levels on LDs. This indicates that decreased LD-associated ATGL is the key factor of insulin resistance. Surprisingly, both PA treatment and ATGL deficiency can upregulate phosphorylated PI3K levels in cells, suggesting that these two factors function within the same signaling pathway. These findings lead us to hypothesize that PA and ATGL may regulate insulin signaling through PIP3 modulation. While the insulin resistance caused by either PA treatment or ATGL deficiency alone can be rescued by elevated phosphorylated PI3K, this rescue mechanism fails when both factors are present, resulting in severe insulin resistance. ATGL deficiency mediated insulin resistance likely involves mechanisms beyond impaired TAG hydrolysis^55,57^, necessitating further investigation to fully elucidate the underlying molecular pathways. Importantly, our results indicate that ATGL deficiency on LDs aggravated PA-induced insulin resistance. In the past, the size and location of skeletal muscle LDs were linked to insulin sensitivity^32^. Type I fibers contain higher LD content than type II fibers, with athletes mainly store lipid in a higher number of LDs in the IMF region of type I fibers, while T2DM patients store lipid in larger LDs in the SS region of type II fibers^34,35^. In the present study, we isolated LDs from total skeletal muscle tissue. It remains technically challenging to isolate LDs from type I and type II muscle fibers separately or accurately distinguish subcellular localization of LDs. Although the regions of LDs in skeletal muscle were not specifically distinguished, the observed differences in the total LD proteome may better reflect physiological variations between groups. Moreover, although our human samples varied in age, sex, and sampling location, we confirmed our key findings through independent experiments in both db/db and ob/ob mouse models. This verification compensates for potential limitations arising from sample heterogeneity and strengthens the reliability of our conclusions.

In conclusion, we isolated and purified skeletal muscle LDs from human participants and performed comparative proteomic analyses of athletes and T2DM patients for the first time. The skeletal muscle LDs in athletes are entirely different from those in T2DM patients. Mitochondrial proteins were upregulated and cytoskeletal proteins were downregulated in the skeletal muscle LDs of athletes. Our results revealed enhanced LDAM in the skeletal muscle of athletes, with LDAM exhibiting elevated oxidative activity. In addition, ATGL was upregulated in the skeletal muscle LDs from athletes, with no changes in whole tissue level. ATGL deficiency aggravated PA-induced insulin resistance. The localization of ATGL on LDs may play an important role in skeletal muscle insulin sensitivity and could represent a potential therapeutic target for alleviating T2DM progression. These findings in the present work support the hypothesis that skeletal muscle contains two distinct types of LDs: the metabolically active, functionally beneficial “good LDs” prevalent in athletes, and the metabolically impaired “bad LDs” characteristic of T2DM patients. Physical interventions such as endurance training may promote the conversion of bad LDs into good LDs, which could significantly contribute to the maintenance of systemic glucose and lipid homeostasis.

## Materials and Methods

### Reagent or Resource

Antibodies against ADRP (Cat#A6276), CGI58 (Cat#A6801), Mettl7B (Cat#A7200), RAB18 (Cat#A2812), and GM130 (Cat#A5344) were purchased from Abclonal. Antibodies against ATGL (Cat#2138S), LAMP1 (Cat#3243), p-AKT (Cat#4051S), and p-PI3K (Cat#4228S) were purchased from Cell Signaling Technology. Antibodies against BiP (Cat#11587-1-AP), AKT (Cat#10176-2-AP), and DHRS1 (Cat#16275-1-AP) were purchased from Proteintech. Antibodies against Sec22B (Cat#SC-101267), Keratin (Cat#SC-376224), and GYS1 (Cat#SC-81173) were purchased from Santa Cruz Biotechnology. LipidTOX Red (Cat#H34476), PageRuler Prestained Protein Ladder (Cat#26617), and Pierce BCA Protein Assay Kit (Cat#23227) were purchased from ThermoFisher Scientific. Insulin was purchased from Beyotime. Sodium palmitate (Cat#P9767) and Sodium oleate (Cat#O7501) were purchased from Sigma-Aldrich. Triacylglycerols Assay Kit (Cat#100000220) was purchased from BioSino Bio-Technology and Science Inc.

### Human Samples

Clinical information and skeletal muscle samples from athletes and T2DM patients were obtained after approval by the Ethical Committee on Human Research of the participating hospitals and with all participants consent. The experiments conformed to the standards set by the Declaration of Helsinki and strictly follow the Ethical Review Measures for Biomedical Research Involving Human Subjects formulated by the National Health Commission of China.

Samples from 7 athletes and 1 T2DM patient were obtained from discarded hamstring muscle tissue during anterior cruciate ligament reconstruction surgery at Peking University Third Hospital. Another 8 samples of T2DM patients were obtained from patients who were admitted to Air Force Medical Center for thoracotomy treatment, these muscle specimens were collected from the pectoralis. A portion of each skeletal muscle sample was removed for LD isolation. The remaining parts of the tissue were fixed in 4% PFA, embedded in paraffin, or flash-frozen in liquid nitrogen for protein analysis. Skeletal muscle morphology and lipid content were evaluated using HE staining and Oil Red O staining. Tissue sections were prepared by Wuhan Servicebio.

### Isolation and Purification of LDs

LDs were purified using a modification of the method used by Liu et al^59^. We collected skeletal muscle samples from athletes or T2DM patients and put them into ice-cold saline containing 0.2 mM PMSF. After removing adipose tissue and fascial tissue, the tissue samples were transferred to moderate volume of Buffer A (250 mM sucrose, 25 mM tricine, pH 7.6) plus 0.2 mM PMSF. The tissue samples were sliced into small fragments by scissors and scalpels, and homogenized with a Tenbroeck tissue grinder (WHEATON, 357424) on ice until complete homogenate formation was achieved. The PNS was obtained by centrifugation of the tissue homogenate at 2,000*g* for 10 min at 4°C. Next, 10 mL PNS was loaded into an SW41 Ti tube and 2 mL Buffer B (20 mM HEPES, pH 7.4, 100 mM KCl, and 2 mM MgCl_2_) was carefully loaded on top. The samples were then centrifuged at 250,000*g* for 1 hour at 4°C. The LDs at the top of the gradients were carefully collected into 1.5 mL EP tubes and washed using Buffer B twice at 20,000*g* for 10 min at 4°C. After washing, Buffer B was removed and a mixture of 1 mL of acetone and 100 µL of chloroform was added to the LDs for lipid extraction and protein precipitation.

### Protein Preparation and Immunoblotting

LD proteins were precipitated by acetone and chloroform at 20,000*g* for 10 min at 4°C. The LD proteins were divided into two aliquots: one was snap-frozen and stored at - 80°C for subsequent mass spectrometry analysis, while the other was dissolved in SDS loading buffer and denatured at 95°C for 5 minutes for silver staining and Western blot analysis. The remaining cellular fraction samples were mixed with an equal volume of 2× SDS sample buffer and stored at 4°C for subsequent analysis. Human tissue samples were prepared according to a previously published method^63^. Briefly, the tissue samples were dissolved in 2× SDS sample buffer and fully homogenized by sonication, followed by protein precipitation using 7.2% trichloroacetic acid (TCA).

Equal amounts of protein lysates were separated by 10% SDS-PAGE, and then either stained with a silver staining kit or transferred to PVDF membranes with 0.45 µm pore size for immunoblotting. The membrane was blocked with 5% fat-free dry milk and incubated overnight at 4°C with the indicated primary antibody. After washing three times for 5 minutes, HRP-conjugated secondary antibody was applied for 1 hour at room temperature. Immunoblotting was detected by ECL system.

### Measurement of TAG Content

The TAG content was measured according to the manufacture’s procedure. Briefly, the tissue samples were treated in saline with 1% Triton X-100 and fully homogenized by sonication. Then, TAG content was determined by TAG assay kit.

### TLC analysis

Lipids from LDs were extracted using a solvent mixture of acetone and chloroform (10:1, v/v). The dried lipids were dissolved in 100 µL chloroform and fractionated by TLC using hexane/ethyl ether/acetic acid (80:20:1, v/v/v). The TLC plate was dried and stained with iodine.

## MS Analysis

Briefly, the protein pellet was dissolved in 20 µL of freshly prepared 8 M urea, reduced with 10 mM DTT at 56°C for 1 hour, and treated with 40 mM iodoacetamide in the dark for 45 minutes to block the sulfhydryl groups. Trypsin was added at a 1:10 (enzyme-to-protein) ratio, followed by overnight digestion at 37°C. After 12 hours, formic acid (FA) was added to quench the reaction. The sample was dried using a speed vacuum and redissolved in FA. The tryptic peptide mixtures were analyzed by LC-MS/MS on an Orbitrap Exploris 480 mass spectrometer. MS/MS data were analyzed using the SEQUEST algorithm against the NCBI RefSeq human protein database. Comparative proteomic analysis with parallel tissue proteome profiling using data-independent acquisition (DIA).

### Confocal Microscopy Observation

Purified LDs with a volume of 10 µL were stained with LipidTOX (1:200) and incubated for 20 minutes. Then, 5 µL of the stained LDs was placed on a glass slide and covered with a coverslip. The LDs were observed using a confocal microscope FV3000.

### Transmission Electron Microscopy Observation

Human skeletal muscle samples were examined with transmission electron microscopy after ultrathin sectioning. Tissues were fixed with 2.5% glutaraldehyde in 0.1 M phosphate buffer (PB, pH 7.4) over night at 4°C, followed by post-fixed in 1% osmium tetraoxide at 4°C for 2 hours. Then the samples were dehydrated through a graded ethanol series at room temperature and then embedded in Embed 812 and subsequently prepared as 70-nm-thick ultrathin sections. Ultrathin sections were examined in an electron microscope. The sections were then observed with Tecnai Spirit electron microscope (FEI, Netherlands) after stained with uranyl acetate and lead citrate.

Isolated LDs suspended in 20 µL Buffer B were mixed with an equal volume of 2% glutaraldehyde in 0.1 M PB and incubated for 30 minutes at room temperature, followed by the addition of 2% OsO₄ and another 30 minutes incubation. Fixed LDs were collected by centrifugation, dehydrated, infiltrated, and embedded in Embed 812. Ultrathin sections were prepared and observed after staining.

For positive staining, LDs were loaded onto Formvar-coated copper grids for 1 minutes, fixed with 2.5% glutaraldehyde for 10 minutes, and 1% OsO₄ for 10 min. Grids were stained with 0.1% tannic acid for 10 minutes and 2% uranyl acetate for 10 minutes, with three 1-minute deionized water washes between steps. Samples were then imaged.

### Cell Culture

C2C12 cells were cultured in Dulbecco’s modified eagle medium (DMEM) supplemented with 10% bovine serum, 100 U/mL penicillin, and 100 μg/mL streptomycin. Cells were incubated at 37°C in a humidified incubator containing 5% CO_2_.

### PA and OA Preparation

PA and OA solutions were prepared according to a previously reported method^64^. Briefly, the fatty acid powder was measured and suspended in ethanol. The suspension was then sonicated to generate a milky emulsion and stored at 4°C for later use.

### Insulin Stimulation

After treating the cells with PA for 20 hours, the culture medium was replaced with fresh medium containing 10 nM insulin, followed by an additional incubation for 15 minutes. Subsequently, the medium was removed, and the cells were lysed with SDS sample buffer for protein sample preparation.

### Animal Studies

ob/ob (C57BLKS/JGpt), ob/ob (C57BL/6JGpt), and their wild-type control mice were purchased from Gempharmatech (Jiangsu, China). All mice were fed with standard rodent chow under a 12 hours light-dark cycle and all experiments were performed when the mice were 8-16 weeks old. All experiments were conducted in compliance with experimental animal ethics guidelines and 3R principles, with proper handling of animals and collection of tissue samples. The experimental procedures were approved by the Animal Care and Use Committee of the Institute of Biophysics, Chinese Academy of Sciences.

### Statistical Analysis

Gene ontology database was used for protein biological process, molecular function and cellular component. KEGG database was used for pathway analysis. These analyses were performed using the DAVID online platform (https://davidbioinformatics.nih.gov/home.jsp). STRING (https://cn.string-db.org) was used for protein interaction. PCA, proteomic clustering heatmap and Venn diagrams were performed using the WeiShengXin (https://www.bioinformatics.com.cn/) online platform. Statistical evaluation of data was performed using two-tailed Student’s t test. *p* < 0.05 was considered as the statistically significant difference. Results were presented as mean ± SD using the software GraphPad Prism.

## Notes

### Competing Interest Statement

The authors have declared no competing interest.

## References

1 Sun, H. et al. IDF Diabetes Atlas: Global, regional and country-level diabetes prevalence estimates for 2021 and projections for 2045. Diabetes Res Clin Pract 183, 109119, doi:10.1016/j.diabres.2021.109119 (2022).

2 Collaborators, G. B. D. D. Global, regional, and national burden of diabetes from 1990 to 2021, with projections of prevalence to 2050: a systematic analysis for the Global Burden of Disease Study 2021. Lancet 402, 203–234, doi:10.1016/S0140-6736(23)01301-6 (2023).

3 Ojo, O. A., Ibrahim, H. S., Rotimi, D. E., Ogunlakin, A. D. & Ojo, A. B. Diabetes mellitus: From molecular mechanism to pathophysiology and pharmacology. Medicine in Novel Technology and Devices 19, doi:10.1016/j.medntd.2023.100247 (2023).

4 DeFronzo, R. A. et al. Type 2 diabetes mellitus. Nat Rev Dis Primers 1, 15019, doi:10.1038/nrdp.2015.19 (2015).

5 Merz, K. E. & Thurmond, D. C. Role of Skeletal Muscle in Insulin Resistance and Glucose Uptake. Compr Physiol 10, 785–809, doi:10.1002/cphy.c190029 (2020).

6 Kahn, S. E., Cooper, M. E. & Del Prato, S. Pathophysiology and treatment of type 2 diabetes: perspectives on the past, present, and future. Lancet 383, 1068–1083, doi:10.1016/S0140-6736(13)62154-6 (2014).

7 Samuel, V. T. & Shulman, G. I. Mechanisms for insulin resistance: common threads and missing links. Cell 148, 852–871, doi:10.1016/j.cell.2012.02.017 (2012).

8 DeFronzo, R. A. & Tripathy, D. Skeletal muscle insulin resistance is the primary defect in type 2 diabetes. Diabetes Care 32 **Suppl 2**, S157–163, doi:10.2337/dc09-S302 (2009).

9 Thiebaud, D. et al. The effect of graded doses of insulin on total glucose uptake, glucose oxidation, and glucose storage in man. Diabetes 31, 957–963, doi:10.2337/diacare.31.11.957 (1982).

10 Ferrannini, E. et al. The disposal of an oral glucose load in patients with non-insulin-dependent diabetes. Metabolism 37, 79–85, doi:10.1016/0026-0495(88)90033-9 (1988).

11 Shulman, G. I. et al. Quantitation of muscle glycogen synthesis in normal subjects and subjects with non-insulin-dependent diabetes by 13C nuclear magnetic resonance spectroscopy. N Engl J Med 322, 223–228, doi:10.1056/NEJM199001253220403 (1990).

12 Liu, P. et al. Chinese hamster ovary K2 cell lipid droplets appear to be metabolic organelles involved in membrane traffic. J Biol Chem 279, 3787–3792, doi:10.1074/jbc.M311945200 (2004).

13 Zhang, S. et al. Morphologically and Functionally Distinct Lipid Droplet Subpopulations. Sci Rep 6, 29539, doi:10.1038/srep29539 (2016).

14 Xu, S., Zhang, X. & Liu, P. Lipid droplet proteins and metabolic diseases. Biochim Biophys Acta Mol Basis Dis 1864, 1968–1983, doi:10.1016/j.bbadis.2017.07.019 (2018).

15 Zhang, C. & Liu, P. The New Face of the Lipid Droplet: Lipid Droplet Proteins. Proteomics 19, e1700223, doi:10.1002/pmic.201700223 (2019).

16 Farese, R. V., Jr. & Walther, T. C. Lipid droplets finally get a little R-E-S-P-E-C-T. Cell 139, 855–860, doi:10.1016/j.cell.2009.11.005 (2009).

17 Olzmann, J. A. & Carvalho, P. Dynamics and functions of lipid droplets. Nat Rev Mol Cell Biol 20, 137–155, doi:10.1038/s41580-018-0085-z (2019).

18 Denton, R. M. & Randle, P. J. Concentrations of glycerides and phospholipids in rat heart and gastrocnemius muscles. Effects of alloxan-diabetes and perfusion. Biochem J 104, 416–422, doi:10.1042/bj1040416 (1967).

19 Damsbo, P., Vaag, A., Hother-Nielsen, O. & Beck-Nielsen, H. Reduced glycogen synthase activity in skeletal muscle from obese patients with and without type 2 (non-insulin-dependent) diabetes mellitus. Diabetologia 34, 239–245, doi:10.1007/BF00405082 (1991).

20 Phillips, D. I. et al. Intramuscular triglyceride and muscle insulin sensitivity: evidence for a relationship in nondiabetic subjects. Metabolism 45, 947–950, doi:10.1016/s0026-0495(96)90260-7 (1996).

21 Pan, D. A. et al. Skeletal muscle triglyceride levels are inversely related to insulin action. Diabetes 46, 983–988, doi:10.2337/diab.46.6.983 (1997).

22 Krssak, M. et al. Intramyocellular lipid concentrations are correlated with insulin sensitivity in humans: a 1H NMR spectroscopy study. Diabetologia 42, 113–116, doi:10.1007/s001250051123 (1999).

23 Manco, M. et al. Insulin resistance directly correlates with increased saturated fatty acids in skeletal muscle triglycerides. Metabolism 49, 220–224, doi:10.1016/s0026-0495(00)91377-5 (2000).

24 van Loon, L. J. Use of intramuscular triacylglycerol as a substrate source during exercise in humans. J Appl Physiol (1985) 97, 1170–1187, doi:10.1152/japplphysiol.00368.2004 (2004).

25 Lee, J. S. et al. Saturated, but not n-6 polyunsaturated, fatty acids induce insulin resistance: role of intramuscular accumulation of lipid metabolites. J Appl Physiol (1985) 100, 1467–1474, doi:10.1152/japplphysiol.01438.2005 (2006).

26 Li, X. et al. Skeletal Muscle Lipid Droplets and the Athlete’s Paradox. Cells 8, doi:10.3390/cells8030249 (2019).

27 Goodpaster, B. H., He, J., Watkins, S. & Kelley, D. E. Skeletal muscle lipid content and insulin resistance: evidence for a paradox in endurance-trained athletes. J Clin Endocrinol Metab 86, 5755–5761, doi:10.1210/jcem.86.12.8075 (2001).

28 Russell, A. P. et al. Lipid peroxidation in skeletal muscle of obese as compared to endurance-trained humans: a case of good vs. bad lipids? FEBS Lett 551, 104–106, doi:10.1016/s0014-5793(03)00875-5 (2003).

29 van Loon, L. J. et al. Intramyocellular lipid content in type 2 diabetes patients compared with overweight sedentary men and highly trained endurance athletes. Am J Physiol Endocrinol Metab 287, E558–565, doi:10.1152/ajpendo.00464.2003 (2004).

30 van Loon, L. J. & Goodpaster, B. H. Increased intramuscular lipid storage in the insulin-resistant and endurance-trained state. Pflugers Arch 451, 606–616, doi:10.1007/s00424-005-1509-0 (2006).

31 Dube, J. J. et al. Exercise-induced alterations in intramyocellular lipids and insulin resistance: the athlete’s paradox revisited. Am J Physiol Endocrinol Metab 294, E882–888, doi:10.1152/ajpendo.00769.2007 (2008).

32 Nielsen, J., Christensen, A. E., Nellemann, B. & Christensen, B. Lipid droplet size and location in human skeletal muscle fibers are associated with insulin sensitivity. Am J Physiol Endocrinol Metab 313, E721–E730, doi:10.1152/ajpendo.00062.2017 (2017).

33 Nielsen, J. et al. Increased subsarcolemmal lipids in type 2 diabetes: effect of training on localization of lipids, mitochondria, and glycogen in sedentary human skeletal muscle. Am J Physiol Endocrinol Metab 298, E706–713, doi:10.1152/ajpendo.00692.2009 (2010).

34 Daemen, S. et al. Distinct lipid droplet characteristics and distribution unmask the apparent contradiction of the athlete’s paradox. Mol Metab 17, 71–81, doi:10.1016/j.molmet.2018.08.004 (2018).

35 Koh, H. E., Nielsen, J., Saltin, B., Holmberg, H. C. & Ortenblad, N. Pronounced limb and fibre type differences in subcellular lipid droplet content and distribution in elite skiers before and after exhaustive exercise. J Physiol 595, 5781–5795, doi:10.1113/JP274462(2017).

36 Hurley, B. F. et al. Muscle triglyceride utilization during exercise: effect of training. J Appl Physiol (1985) 60, 562–567, doi:10.1152/jappl.1986.60.2.562 (1986).

37 Tarnopolsky, M. A. et al. Influence of endurance exercise training and sex on intramyocellular lipid and mitochondrial ultrastructure, substrate use, and mitochondrial enzyme activity. Am J Physiol Regul Integr Comp Physiol 292, R1271–1278, doi:10.1152/ajpregu.00472.2006 (2007).

38 Cui, L., Mirza, A. H., Zhang, S., Liang, B. & Liu, P. Lipid droplets and mitochondria are anchored during brown adipocyte differentiation. Protein Cell 10, 921–926, doi:10.1007/s13238-019-00661-1 (2019).

39 Yu, J. et al. Lipid droplet remodeling and interaction with mitochondria in mouse brown adipose tissue during cold treatment. Biochim Biophys Acta 1853, 918–928, doi:10.1016/j.bbamcr.2015.01.020 (2015).

40 Pu, J. et al. Interactomic study on interaction between lipid droplets and mitochondria. Protein Cell 2, 487–496, doi:10.1007/s13238-011-1061-y (2011).

41 Wang, H. et al. Perilipin 5, a lipid droplet-associated protein, provides physical and metabolic linkage to mitochondria. J Lipid Res 52, 2159–2168, doi:10.1194/jlr.M017939 (2011).

42 Cui, L. & Liu, P. Two Types of Contact Between Lipid Droplets and Mitochondria. Front Cell Dev Biol 8, 618322, doi:10.3389/fcell.2020.618322 (2020).

43 Zhang, H. et al. Proteome of skeletal muscle lipid droplet reveals association with mitochondria and apolipoprotein a-I. J Proteome Res 10, 4757–4768, doi:10.1021/pr200553c (2011).

44 Li, L. et al. Comparative proteomics reveals abnormal binding of ATGL and dysferlin on lipid droplets from pressure overload-induced dysfunctional rat hearts. Sci Rep 6, 19782, doi:10.1038/srep19782 (2016).

45 Nielsen, J. et al. Increased contact between lipid droplets and mitochondria in skeletal muscles of male elite endurance athletes. Am J Physiol Cell Physiol, doi:10.1152/ajpcell.00123.2025 (2025).

46 Renne, M. F. & Hariri, H. Lipid Droplet-Organelle Contact Sites as Hubs for Fatty Acid Metabolism, Trafficking, and Metabolic Channeling. Front Cell Dev Biol 9, 726261, doi:10.3389/fcell.2021.726261 (2021).

47 Yang, M. et al. Lipid droplet - mitochondria coupling: A novel lipid metabolism regulatory hub in diabetic nephropathy. Front Endocrinol (Lausanne) 13, 1017387, doi:10.3389/fendo.2022.1017387 (2022).

48 Enkler, L. & Spang, A. Functional interplay of lipid droplets and mitochondria. FEBS Lett 598, 1235–1251, doi:10.1002/1873-3468.14809 (2024).

49 Fan, H. & Tan, Y. Lipid Droplet-Mitochondria Contacts in Health and Disease. Int J Mol Sci 25, doi:10.3390/ijms25136878 (2024).

50 Smolkova, K. & Gotvaldova, K. Fatty Acid Trafficking Between Lipid Droplets and Mitochondria: An Emerging Perspective. Int J Biol Sci 21, 1863–1873, doi:10.7150/ijbs.105361 (2025).

51 Czajkowska, A. et al. Exploring protein relative relations in skeletal muscle proteomic analysis for insights into insulin resistance and type 2 diabetes. Sci Rep 14, 17631, doi:10.1038/s41598-024-68568-4 (2024).

52 Kjaergaard, J. et al. Personalized molecular signatures of insulin resistance and type 2 diabetes. Cell, doi:10.1016/j.cell.2025.05.005 (2025).

53 Mulder, H. et al. Hormone-sensitive lipase null mice exhibit signs of impaired insulin sensitivity whereas insulin secretion is intact. J Biol Chem 278, 36380–36388, doi:10.1074/jbc.M213032200 (2003).

54 Park, S. Y. et al. Hormone-sensitive lipase knockout mice have increased hepatic insulin sensitivity and are protected from short-term diet-induced insulin resistance in skeletal muscle and heart. Am J Physiol Endocrinol Metab 289, E30–39, doi:10.1152/ajpendo.00251.2004 (2005).

55 Haemmerle, G. et al. Defective lipolysis and altered energy metabolism in mice lacking adipose triglyceride lipase. Science 312, 734–737, doi:10.1126/science.1123965 (2006).

56 Schoenborn, V. et al. The ATGL gene is associated with free fatty acids, triglycerides, and type 2 diabetes. Diabetes 55, 1270–1275, doi:10.2337/db05-1498 (2006).

57 Kienesberger, P. C. et al. Adipose triglyceride lipase deficiency causes tissue-specific changes in insulin signaling. J Biol Chem 284, 30218–30229, doi:10.1074/jbc.M109.047787 (2009).

58 Albert, J. S. et al. Null mutation in hormone-sensitive lipase gene and risk of type 2 diabetes. N Engl J Med 370, 2307–2315, doi:10.1056/NEJMoa1315496 (2014).

59 Ding, Y. et al. Isolating lipid droplets from multiple species. Nat Protoc 8, 43–51, doi:10.1038/nprot.2012.142 (2013).

60 Chen, X. et al. Comparative Proteomic Study of Fatty Acid-treated Myoblasts Reveals Role of Cox-2 in Palmitate-induced Insulin Resistance. Sci Rep 6, 21454, doi:10.1038/srep21454 (2016).

61 Peng, G. et al. Oleate blocks palmitate-induced abnormal lipid distribution, endoplasmic reticulum expansion and stress, and insulin resistance in skeletal muscle. Endocrinology 152, 2206–2218, doi:10.1210/en.2010-1369 (2011).

62 Pu, J. et al. Palmitic acid acutely stimulates glucose uptake via activation of Akt and ERK1/2 in skeletal muscle cells. J Lipid Res 52, 1319–1327, doi:10.1194/jlr.M011254 (2011).

63 Cheng, L. et al. Protein sample preparation for tissue distribution study. Proteomics Clin Appl 17, e2200088, doi:10.1002/prca.202200088 (2023).

64 Zhang, S. et al. Preparation of fatty acid solutions for investigating lipid signaling, metabolism, and lipid droplets. Protein Cell, doi:10.1093/procel/pwae068 (2024).

