## Supplemental Data for "Lipid Droplet Proteome Reveals that Associated ATGL and Anchored Mitochondria Lead to Higher Skeletal Muscle Insulin Sensitivity in Endurance Athletes than Type 2 Diabetes Mellitus"

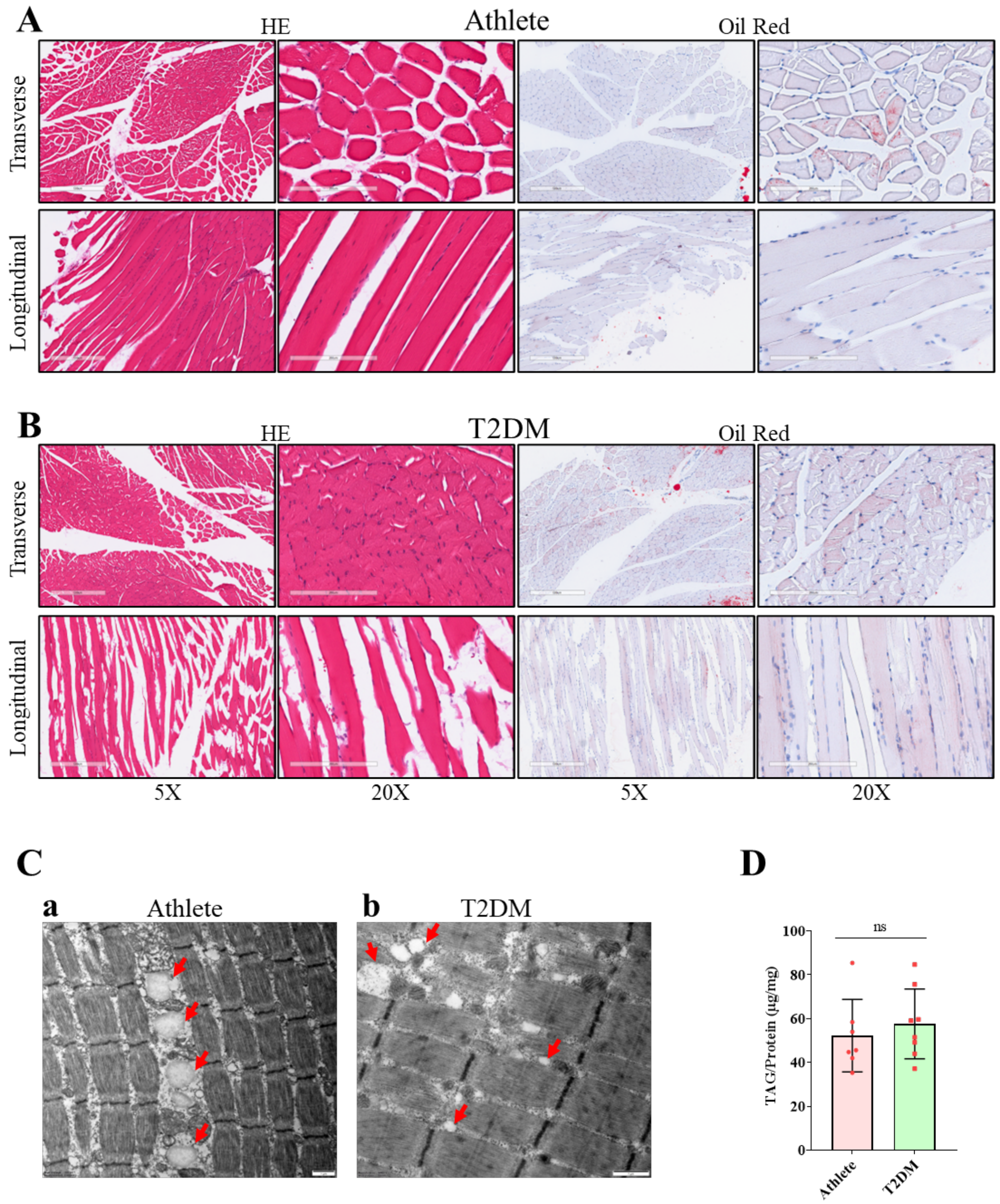


**Figure S1 Histomorphometric and biochemical analyses of intramuscular lipid deposition in athletes and T2DM patients.**

(A) Observation of the morphological characteristics by HE staining and the lipid profile by Oil Red O staining in skeletal muscle from athletes. Muscle tissue sections were prepared in both transverse and longitudinal orientations, with representative images captured at 5× and 20× magnification. Scale bars represent 200 μm (20× magnification) and 500 μm (5× magnification).

(B) Observation of the morphological characteristics by HE staining and the lipid profile by Oil Red O staining in skeletal muscle from T2DM patients. Muscle tissue sections were prepared in both transverse and longitudinal orientations, with representative images captured at 5× and 20× magnification. Scale bars represent 200 μm (20× magnification) and 500 μm (5× magnification).

(C) Morphology of skeletal muscle LDs in tissues from athletes (a) and T2DM patients (b). Skeletal muscle was sectioned to 70 nm thick, stained with 2% uranyl acetate and observed by TEM. LDs are indicated by red arrowheads. Scale bar = 1 μm.

(D) TAG content in skeletal muscle tissues from athletes and T2DM patients.

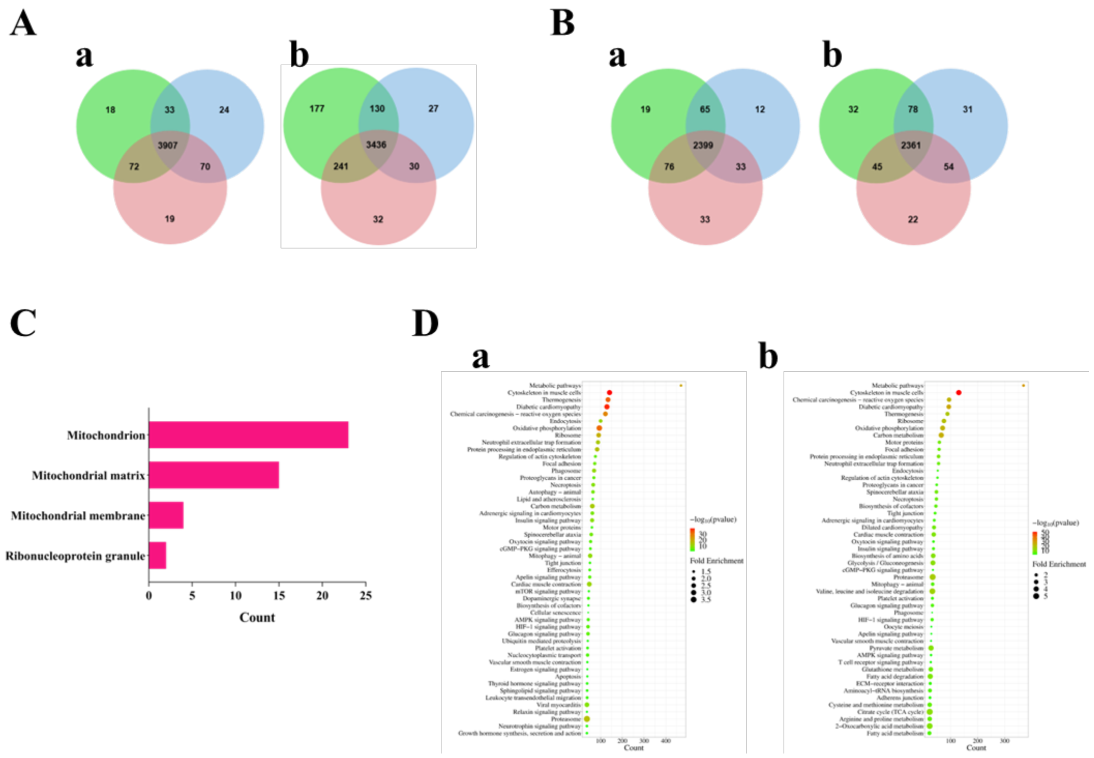


**Figure S2 Comparative proteomic analysis of LDs and tissues from human participants.**

(A) Venn diagram shows the overlap of identified LD proteins between 3 athletes (a) and 3 T2DM patients (b).

(B) Venn diagram shows the overlap of identified tissue proteins between 3 athletes (a) and 3 T2DM patients (b).

(C) GOCC enrichment analysis of proteins only present in 3 LD samples from athletes.

(D) KEGG pathway enrichment analysis was performed on proteins identified with unique peptides ≥ 3 across all six LD samples (a) or six tissue samples (b).

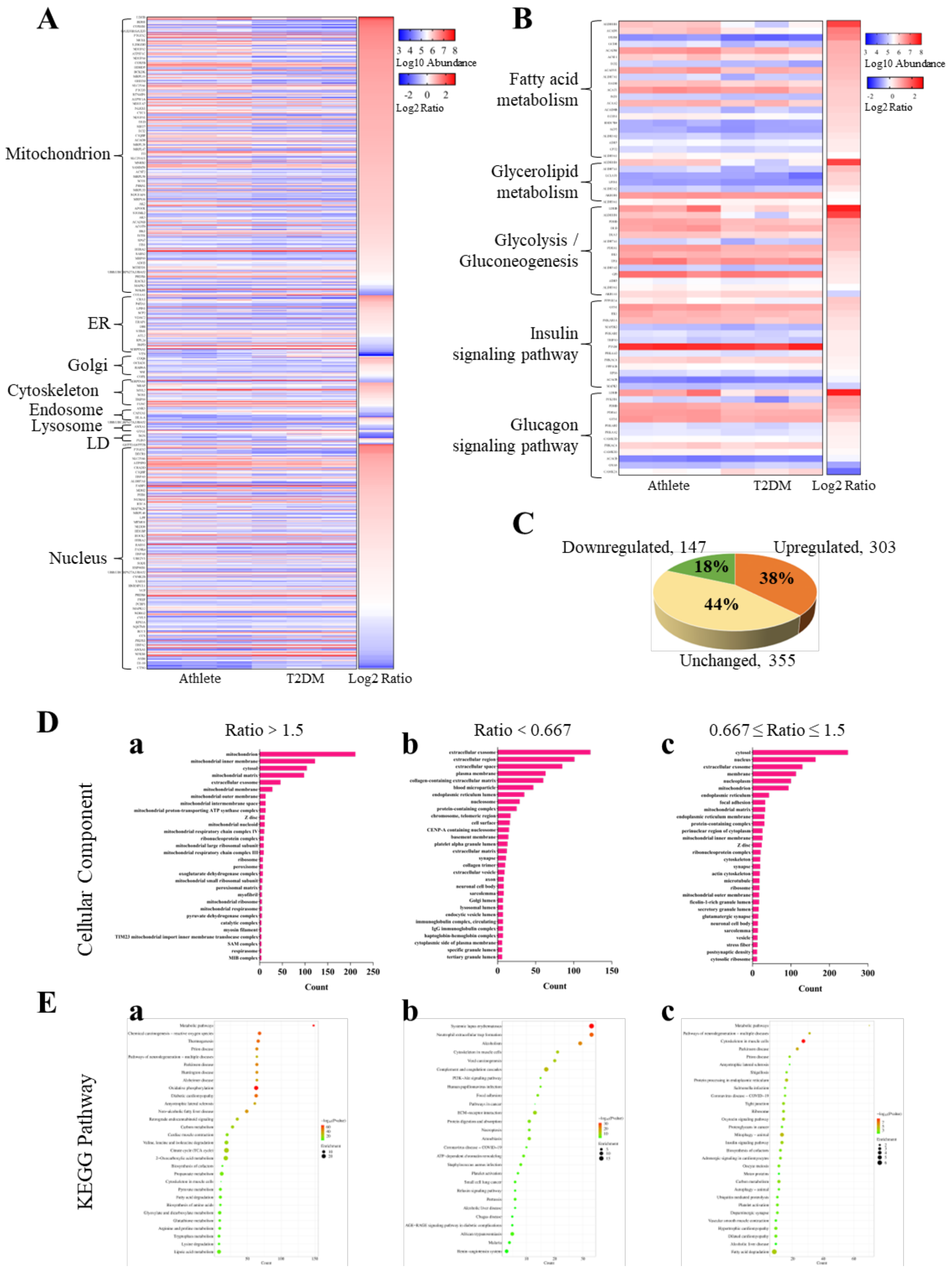


**Figure S3 Comparative proteomic analysis of tissues between athletes and T2DM patients.**

(A) The heatmap of differentially expressed proteins in the tissue proteome. Proteins with unique peptide ≥ 3 and $p < 0.05$ were selected for analysis by GOCC enrichment. The color gradient represents the log10-transformed abundance of each protein, while the ratio color intensity reflects the log2-transformed fold change (athletes vs. T2DM patients).

(B) The heatmap of differentially expressed functional pathway proteins in the tissue proteome. Proteins with unique peptide $\geq 3$ and $p < 0.05$ were initially selected by KEGG pathway enrichment and subsequently filtered to specifically identify those involved in fatty acid metabolism, glycerolipid metabolism, glycolysis/gluconeogenesis, insulin signaling pathway, and glucagon signaling pathway for further analysis.

(C) Percentages of upregulated, downregulated, and unchanged proteins relative to the total protein ($p < 0.05$) in the tissue proteome.

(D) Top 30 GOCC enrichment categories for upregulated (Ratio > 1.5, a), downregulated (Ratio < 0.667, b), and unchanged (0.667 ≤ Ratio ≤ 1.5, c) proteins identified in the LD proteome.

(E) Top 30 KEGG enrichment pathways for upregulated (Ratio > 1.5, a), downregulated (Ratio < 0.667, b), and unchanged (0.667 ≤ Ratio ≤ 1.5, c) proteins identified in the LD proteome.

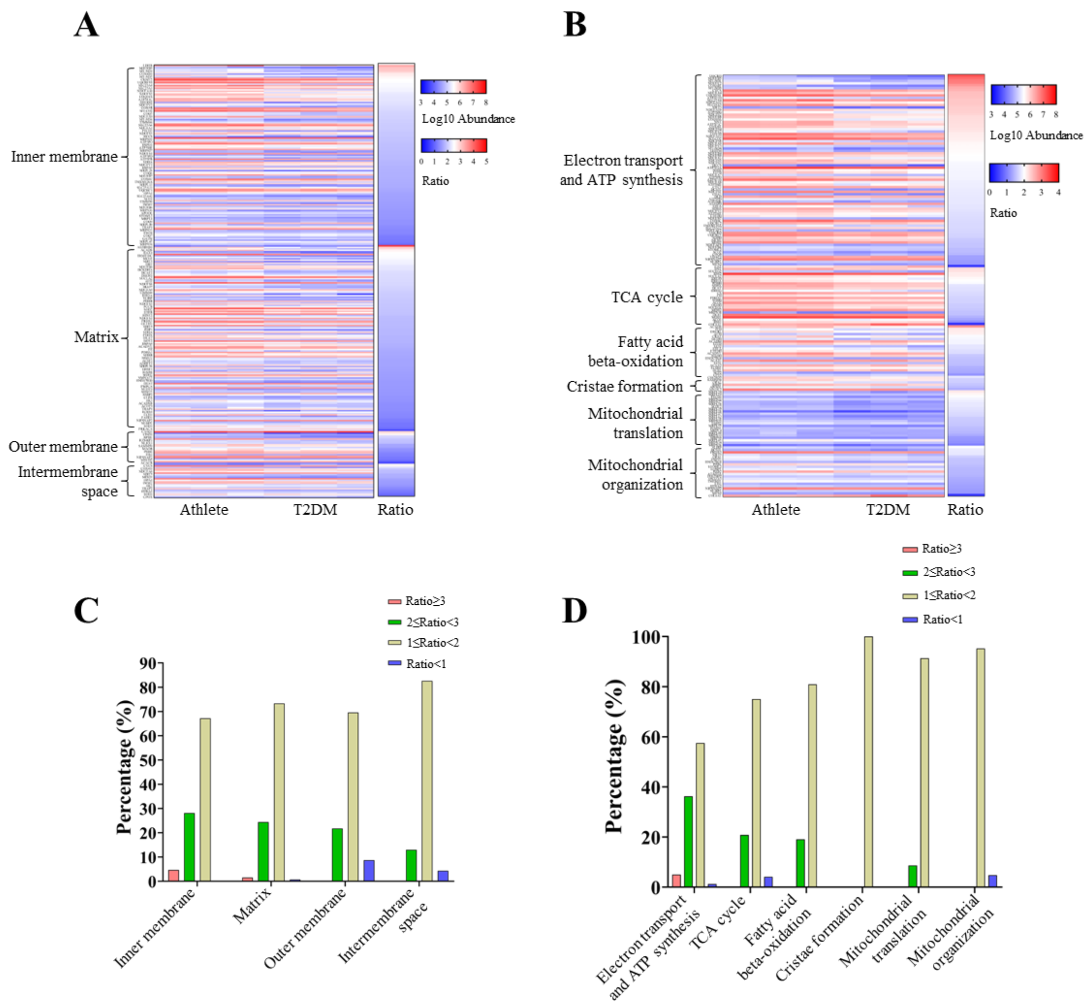


**Figure S4 Increased mitochondria in the skeletal muscle tissues of athletes.**

(A) The heatmap of differentially expressed mitochondrial subcompartment proteins in the tissue proteome. Mitochondrial proteins with unique peptide ≥ 3 and $p < 0.05$ were selected for analysis by GOCC enrichment.

(B) The heatmap of differentially expressed mitochondrial functional proteins in the tissue proteome. Mitochondrial proteins with unique peptide ≥ 3 and $p < 0.05$ were initially selected by GOCC enrichment and subsequently filtered through GOBP enrichment for further analysis.

(C) The percentage distribution of abundance ratios for mitochondrial proteins across the four mitochondrial subcompartments in the tissue proteome. The data is derived from the results in figure A.

(D) The percentage distribution of abundance ratios for mitochondrial functional proteins in the tissue proteome. The data is derived from the results in figure B.

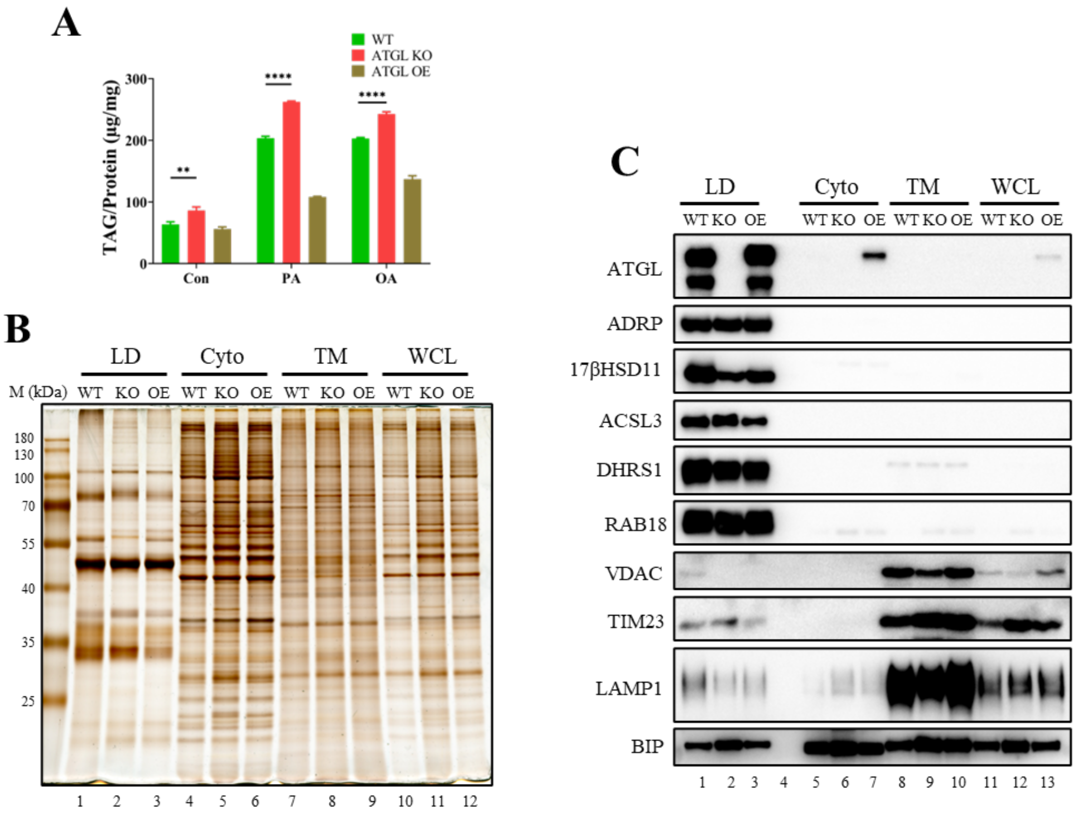


**Figure S5 Isolation and characterization of LDs from ATGL knockout and overexpression cells.**

(A) Measurement of TAG content in ATGL KO and OE cells. PA was used at a concentration of 200 μM, OA at 100 μM.

(B) Protein fractions isolated from ATGL KO and OE cells, including lipid droplets (LDs), cytosol (Cyto), total membrane (TM), and whole cell lysate (WCL), were subjected to silver staining. The cells were treated by 100 μM OA for 12 hours before LD isolation.

(C) Protein fractions isolated from ATGL KO and OE cells were analyzed by Western blot.

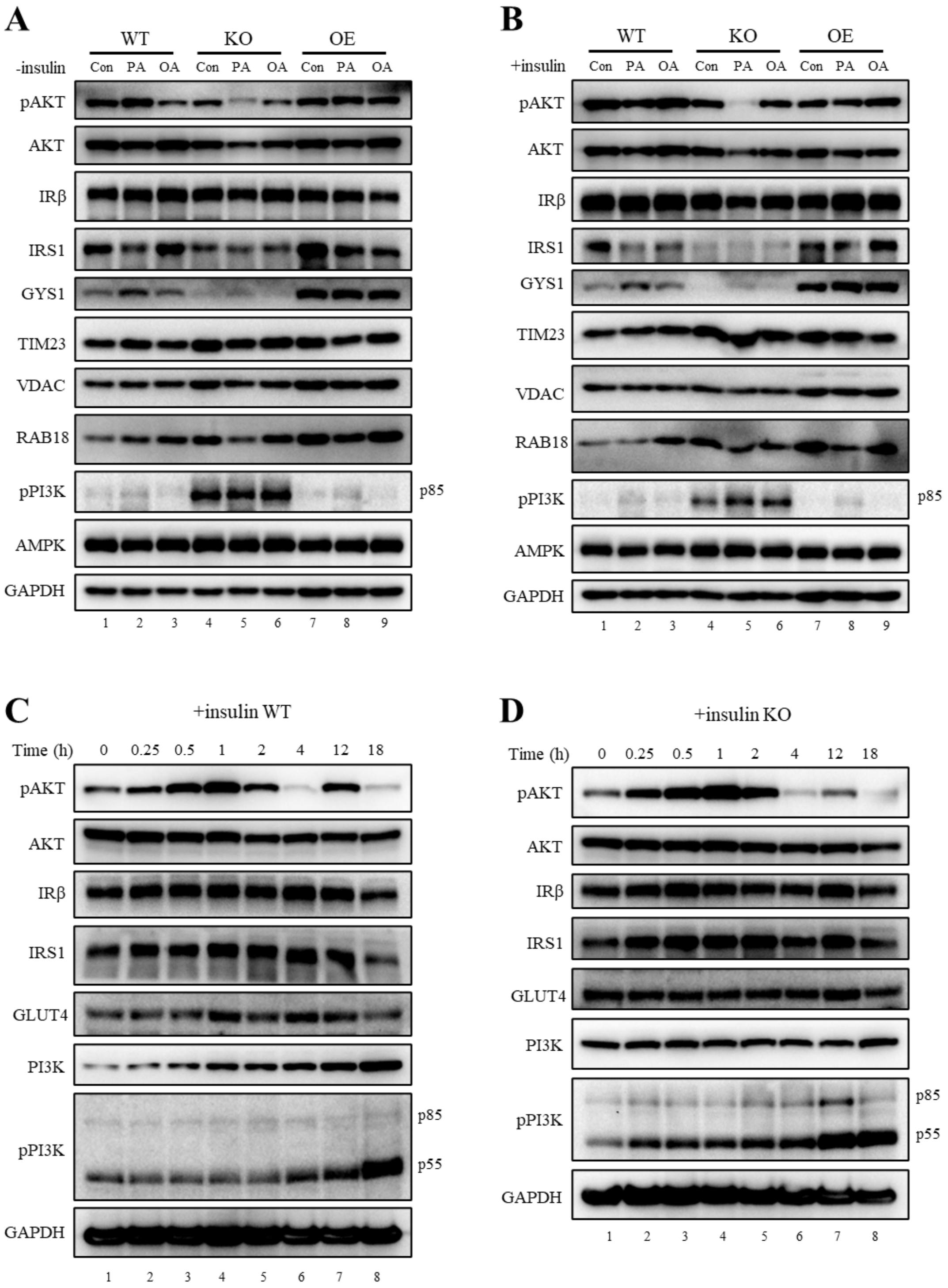


**Figure S6 Impact of insulin addition and PA treatment timing on AKT phosphorylation induced by ATGL deficiency.**

(A) Effect of ATGL knockout on AKT phosphorylation in the absence of insulin supplementation in culture media. Cells were sampled directly after PA treatment and then detected by immunoblotting.

(B) Effect of ATGL knockout on AKT phosphorylation in the presence of insulin supplementation in culture media. After treatment with PA, cells were further incubated with 10 nM insulin for 15 minutes, followed by sample preparation for Western blot analysis.

(C) Time course of PA treatment on AKT phosphorylation dynamics in WT C2C12 cells.

(D) Time course of PA treatment on AKT phosphorylation dynamics in ATGL KO cells.

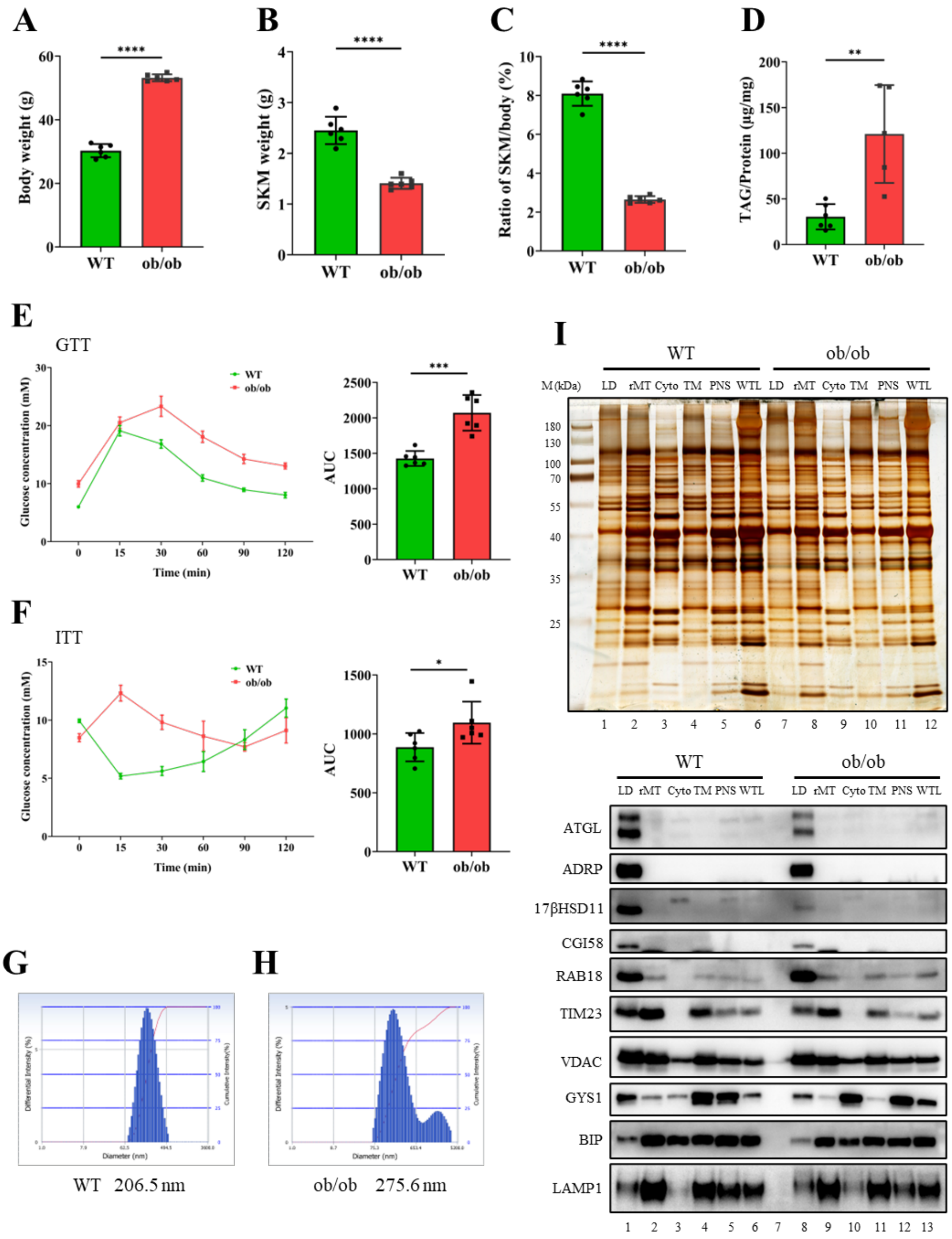


**Figure S7 ATGL reduction and mitochondrial protein increase in skeletal muscle LDs of ob/ob mice.**

(A) Body weight of WT and ob/ob mice. ****, $p < 0.0001$.

(B) Skeletal muscle weight from four limbs in WT and ob/ob mice after removal of adipose tissue. ****, $p < 0.0001$.

(C) Ratio of skeletal muscle mass from four limbs to body weight in mice. ****, $p <$

0.0001.

(D) TAG content of skeletal muscle tissues in WT and ob/ob mice. **, $p < 0.01$.

(E) Glucose tolerance test in ob/ob mice. Following 16 hours of fasting, glucose was injected intraperitoneally to mice (1.25 g/kg body weight). ***, $p < 0.001$.

(F) Insulin tolerance test in ob/ob mice. Following 3 hours of fasting, insulin was injected intraperitoneally to mice (0.5 IU/kg body weight). *, $p < 0.05$.

(G) The size of LDs from skeletal muscle in WT mice. Skeletal muscles from four limbs of three WT mice were mixed for LD isolation.

(H) The size of LDs from skeletal muscle in ob/ob mice. Skeletal muscles from four limbs of three ob/ob mice were mixed for LD isolation.

(I) Protein fractions isolated from skeletal muscle, including lipid droplets (LDs), rough mitochondria (rMT), cytosol (Cyto), total membrane (TM), postnuclear supernatant (PNS) and whole tissue lysate (WTL), were subjected to silver staining and further immunoblot analysis. Protein loading normalization based on equivalent TAG content for LD samples. ATGL expression along with other proteins of interest (ADRP, TIM23, BIP, and LAMP1) was assessed by Western blot analysis.

**Table S1 The top 50 upregulated proteins in LDs from skeletal muscle of athletes compared with T2DM.**

| Genes | Protein Descriptions | Unique Peptides | Ratio | Q value |
|---|---|---|---|---|
| **ALDH1B1** | Aldehyde dehydrogenase X, mitochondrial | 21 | 12.136 | 0.0041 |
| **ACSF3** | Malonate--CoA ligase ACSF3, mitochondrial | 10 | 7.787 | 0.0134 |
| **MRPL47** | 39S ribosomal protein L47, mitochondrial | 8 | 6.780 | 0.0007 |
| **CARS2** | Probable cysteine--tRNA ligase, mitochondrial | 14 | 6.045 | 0.0039 |
| **DNAJC14** | DnaJ homolog subfamily C member 14 | 7 | 5.953 | 0.0433 |
| **DHRS1** | Dehydrogenase/reductase SDR family member 1 | 7 | 5.307 | 0.0029 |
| **VCP** | Transitional endoplasmic reticulum ATPase | 43 | 4.928 | 0.0082 |
| **ATP5IF1** | ATPase inhibitor, mitochondrial | 14 | 4.922 | 0.0008 |
| **GATD3** | Glutamine amidotransferase-like class 1 domain-containing protein 3, mitochondrial | 14 | 4.778 | 0.0100 |
| **LDHB** | L-lactate dehydrogenase B chain | 9 | 4.529 | 0.0048 |
| **TRMT10C** | tRNA methyltransferase 10 homolog C | 13 | 4.472 | 0.0282 |
| **G0S2** | G0/G1 switch protein 2 | 6 | 4.371 | 0.0024 |
| **PPIF** | Peptidyl-prolyl cis-trans isomerase F, mitochondrial | 7 | 4.277 | 0.0077 |
| **NCAM1** | Neural cell adhesion molecule 1 | 19 | 4.256 | 0.0062 |
| **PPOX** | Protoporphyrinogen oxidase | 12 | 4.155 | 0.0008 |
| **NNT** | NAD(P) transhydrogenase, mitochondrial | 68 | 4.035 | 0.0026 |
| **COX6A2** | Cytochrome c oxidase subunit 6A2, mitochondrial | 6 | 4.005 | 0.0001 |
| **ACADS** | Short-chain specific acyl-CoA dehydrogenase, mitochondrial | 18 | 3.976 | 0.0019 |
| **ABCB7** | Iron-sulfur clusters transporter ABCB7, mitochondrial | 14 | 3.971 | 0.0012 |
| **DLAT** | Dihydrolipoyllysine-residue acetyltransferase component of pyruvate dehydrogenase complex, mitochondrial | 33 | 3.890 | 0.0008 |
| **SLC25A4** | ADP/ATP translocase 1 | 22 | 3.842 | 0.0008 |
| **C6orf136** | Uncharacterized protein C6orf136 | 5 | 3.816 | 0.0003 |
| **GYS1** | Glycogen [starch] synthase, muscle | 38 | 3.793 | 0.0000 |
| **LDAH** | Lipid droplet-associated hydrolase | 6 | 3.741 | 0.0010 |
| **BCS1L** | Mitochondrial chaperone BCS1 | 16 | 3.730 | 0.0015 |
| **MRPS36** | Alpha-ketoglutarate dehydrogenase component 4 | 10 | 3.584 | 0.0004 |
| **ACADSB** | Short/branched chain specific acyl-CoA dehydrogenase, mitochondrial | 16 | 3.573 | 0.0042 |
| **GCDH** | Glutaryl-CoA dehydrogenase, mitochondrial | 10 | 3.559 | 0.0013 |
| **PC** | Pyruvate carboxylase, mitochondrial | 18 | 3.543 | 0.0010 |
| **TUFM** | Elongation factor Tu, mitochondrial | 29 | 3.532 | 0.0044 |
| **PTRHD1** | Putative peptidyl-tRNA hydrolase PTRHD1 | 5 | 3.517 | 0.0028 |
| **GADD45GIP1** | Growth arrest and DNA damage-inducible proteins-interacting protein 1 | 6 | 3.506 | 0.0038 |
| **SLC25A25** | Calcium-binding mitochondrial carrier protein SCaMC-2 | 3 | 3.505 | 0.0091 |
| **MRPL13** | 39S ribosomal protein L13, mitochondrial | 11 | 3.407 | 0.0016 |
| **PRODH** | Proline dehydrogenase 1, mitochondrial | 12 | 3.384 | 0.0090 |
| **PDHA1** | Pyruvate dehydrogenase E1 component subunit alpha, somatic form, mitochondrial | 35 | 3.378 | 0.0007 |

| | | | | |
|---|---|---|---|---|
| **ISOC2** | Isochorismatase domain-containing protein 2 | 5 | 3.376 | 0.0081 |
| **ACADVL** | Very long-chain specific acyl-CoA dehydrogenase, mitochondrial | 91 | 3.376 | 0.0011 |
| **COQ10A** | Coenzyme Q-binding protein COQ10 homolog A, mitochondrial | 8 | 3.361 | 0.0011 |
| **SOD2** | Superoxide dismutase [Mn], mitochondrial | 15 | 3.355 | 0.0037 |
| **IDH3B** | Isocitrate dehydrogenase [NAD] subunit beta, mitochondrial | 16 | 3.330 | 0.0029 |
| **DLD** | Dihydrolipoyl dehydrogenase, mitochondrial | 34 | 3.319 | 0.0003 |
| **HSPE1** | 10 kDa heat shock protein, mitochondrial | 10 | 3.316 | 0.0149 |
| **COMTD1** | Catechol O-methyltransferase domain-containing protein 1 | 7 | 3.315 | 0.0082 |
| **METTL7B** | Thiol S-methyltransferase METTL7B | 7 | 3.311 | 0.0268 |
| **SPATA20** | Spermatogenesis-associated protein 20 | 12 | 3.287 | 0.0138 |
| **MCCC1** | Methylcrotonoyl-CoA carboxylase subunit alpha, mitochondrial | 14 | 3.284 | 0.0042 |
| **ACOT1** | Acyl-coenzyme A thioesterase 1 | 21 | 3.280 | 0.0129 |
| **MRPL2** | 39S ribosomal protein L2, mitochondrial | 4 | 3.267 | 0.0009 |
| **ATP2A2** | Sarcoplasmic/endoplasmic reticulum calcium ATPase 2 | 52 | 3.261 | 0.0036 |

**Table S2 The top 50 downregulated proteins in LDs from skeletal muscle of athletes compared with T2DM.**

| Genes | Protein Descriptions | Unique Peptides | Ratio | Q value |
|---|---|---|---|---|
| **KRT16** | Keratin, type I cytoskeletal 16 | 18 | 0.026 | 0.0065 |
| **CAMP** | Cathelicidin antimicrobial peptide | 3 | 0.035 | 0.0043 |
| **TMOD3** | Tropomodulin-3 | 3 | 0.041 | 0.0054 |
| **KRT17** | Keratin, type I cytoskeletal 17 | 10 | 0.046 | 0.0081 |
| **NEXN** | Nexilin | 24 | 0.047 | 0.0059 |
| **HBD** | Hemoglobin subunit delta | 5 | 0.050 | 0.0028 |
| **HBQ1** | Hemoglobin subunit theta-1 | 4 | 0.051 | 0.0050 |
| **SERPIND1** | Heparin cofactor 2 | 7 | 0.052 | 0.0005 |
| **S100A8** | Protein S100-A8 | 4 | 0.059 | 0.0010 |
| **CA1** | Carbonic anhydrase 1 | 14 | 0.066 | 0.0032 |
| **COMMD8** | COMM domain-containing protein 8 | 3 | 0.066 | 0.0324 |
| **APCS** | Serum amyloid P-component | 7 | 0.067 | 0.0016 |
| **FGA** | Fibrinogen alpha chain | 22 | 0.068 | 0.0048 |
| **C4BPA** | C4b-binding protein alpha chain | 9 | 0.069 | 0.0016 |
| **FGB** | Fibrinogen beta chain | 18 | 0.070 | 0.0044 |
| **ITGA2B** | Integrin alpha-IIb | 14 | 0.070 | 0.0061 |
| **SYNPO2** | Synaptopodin-2 | 30 | 0.071 | 0.0112 |
| **DDX28** | Probable ATP-dependent RNA helicase DDX28 | 10 | 0.075 | 0.0492 |
| **CTSG** | Cathepsin G | 4 | 0.076 | 0.0073 |
| **COL1A1** | Collagen alpha-1(I) chain | 24 | 0.076 | 0.0031 |
| **APOC4** | Apolipoprotein C-IV | 5 | 0.078 | 0.0021 |
| **TMOD1** | Tropomodulin-1 | 18 | 0.081 | 0.0025 |
| **COL1A2** | Collagen alpha-2(I) chain | 14 | 0.082 | 0.0030 |
| **KRT6B** | Keratin, type II cytoskeletal 6B | 6 | 0.083 | 0.0091 |
| **SAA1** | Serum amyloid A-1 protein | 8 | 0.084 | 0.0088 |
| **XIRP1** | Xin actin-binding repeat-containing protein 1 | 53 | 0.085 | 0.0086 |
| **PROS1** | Vitamin K-dependent protein S | 10 | 0.085 | 0.0048 |
| **SYNE1** | Nesprin-1 | 4 | 0.087 | 0.0237 |
| **APOC2** | Apolipoprotein C-II | 5 | 0.088 | 0.0001 |
| **ACTN3** | Alpha-actinin-3 | 37 | 0.088 | 0.0102 |
| **C4A** | Complement C4-A | 33 | 0.089 | 0.0012 |
| **TNNC2** | Troponin C, skeletal muscle | 15 | 0.091 | 0.0036 |
| **COL3A1** | Collagen alpha-1(III) chain | 7 | 0.091 | 0.0024 |
| **KRT14** | Keratin, type I cytoskeletal 14 | 30 | 0.095 | 0.0074 |
| **HBB** | Hemoglobin subunit beta | 18 | 0.095 | 0.0027 |
| **HBZ** | Hemoglobin subunit zeta | 7 | 0.095 | 0.0098 |
| **HTT** | Huntingtin | 18 | 0.096 | 0.0193 |
| **CAPZA1** | F-actin-capping protein subunit alpha-1 | 6 | 0.098 | 0.0053 |
| **MYOZ1** | Myozenin-1 | 16 | 0.099 | 0.0057 |
| **KRT6A** | Keratin, type II cytoskeletal 6A | 33 | 0.100 | 0.0047 |
| **LUM** | Lumican | 11 | 0.102 | 0.0102 |

| | | | | |
|---|---|---|---|---|
| **FGG** | Fibrinogen gamma chain | 17 | 0.105 | 0.0068 |
| **TNNT3** | Troponin T, fast skeletal muscle | 26 | 0.107 | 0.0038 |
| **CAPZA2** | F-actin-capping protein subunit alpha-2 | 11 | 0.109 | 0.0088 |
| **SYNPO2L** | Synaptopodin 2-like protein | 21 | 0.109 | 0.0100 |
| **COL4A2** | Collagen alpha-2(IV) chain | 7 | 0.109 | 0.0030 |
| **TPM1** | Tropomyosin alpha-1 chain | 22 | 0.111 | 0.0044 |
| **DSC1** | Desmocollin-1 | 8 | 0.115 | 0.0003 |
| **IGHA1** | Immunoglobulin heavy constant alpha 1 | 7 | 0.115 | 0.0027 |
| **HBG1** | Hemoglobin subunit gamma-1 | 10 | 0.116 | 0.0066 |
